# Gdnf, a germ cell-derived factor, regulates zebrafish germ cell stemness through the creation of new spermatogonial niches (germ and Sertoli cells) and inhibition of spermatogonial differentiation in an autocrine and paracrine manners

**DOI:** 10.1101/2021.12.25.474164

**Authors:** Lucas B. Doretto, Arno J. Butzge, Rafael T. Nakajima, Emanuel R. M. Martinez, Beatriz Marques, Maira da Silva Rodrigues, Ivana F. Rosa, Juliana M. B. Ricci, Aldo Tovo-Neto, Daniel F. Costa, Guilherme Malafaia, Changwei Shao, Rafael H. Nóbrega

## Abstract

Glial cell line-derived neurotrophic factor (GDNF) and its receptor (GDNF Family Receptor α1 - GFRα1) are well known to mediate spermatogonial stem cell (SSC) proliferation and survival in the mammalian testes. In nonmammalian species, Gdnf and Gfrα1 orthologs have been found but their functions remain poorly investigated in the testis. Considering this background, this study aimed to understand the roles of Gdnf-Gfrα1 signaling pathway in the zebrafish testis by combining *in vivo*, *in silico* and *ex vivo* approaches. Our analysis showed that zebrafish exhibited two paralogs of Gndf (*gdnfa* and *gdnfb*) and its receptor, Gfrα1 (*gfrα1a* and *gfrα1b*), in agreement with the teleost-specific third round (3R) of whole genome duplication. Expression analysis further revealed that *gdnfa* and *gfrα1a* were the most expressed copies in the zebrafish adult testes. Subsequently, we demonstrated that *gdnfa* is expressed in the germ cells, while Gfrα1a was detected in early spermatogonia (mainly in types A_und_ and A_diff_) and Sertoli cells. Functional *ex vivo* analysis showed that Gdnf promoted the creation of new available niches by stimulating proliferation of both type A_und_ spermatogonia and their surrounding Sertoli cells, but without changing *pou5f3* mRNA levels. Strikingly, Gdnf also inhibited late spermatogonial differentiation as shown by the decrease of type B spermatogonia and down-regulation of *dazl* in the co-treatment with Fsh. Altogether, our data revealed for the first time that a germ cell-derived factor is associated with maintaining germ cell stemness through the creation of new available niches, supporting development of differentiating spermatogonial cysts and inhibiting late spermatogonial differentiation in autocrine and paracrine manners.

## Introduction

GDNF (Glial cell line-derived neurotrophic factor) is a closely related member of TGF-β superfamily which belongs to the GDNF family of ligands (GFLs). This family of ligands consists of Gdnf, neurturin, artemin and persephin [1]. The importance of GDNF for spermatogonial stem cell (SSC) maintenance was unveiled by Meng et al. [2] who showed that mice with impaired GDNF signaling exhibited a progressive loss of SSCs, while its pan-ectopic overexpression promoted germ cell hyperplasia, and ultimately tumors [2]. Moreover, further studies showed that GDNF promoted *in vitro* expansion of mouse germline stem cells [3, 4], being considered as an indispensable factor for long-term culture of SSCs of several rodents [3, 5, 6]. More recently, experiments using mice that ectopically expressed stage-specific GDNF in Sertoli cells revealed that GDNF increased SSC self-renewal by blocking differentiation rather than actively stimulating their proliferation [7]. Altogether, these studies demonstrated that GDNF is an important factor for mammalian SSC self-renewal, proliferation of the stem cell direct progenitors and maintenance of the SSC undifferentiated state (see review in Parekh et al. [8] Mäkelä and Hobbs [9]).

GDNF signaling occurs through binding the non-signaling co-receptor of the GDNF Family Receptor α1 (GFRα1), which are tethered to the plasma membrane through glycosylphosphatidylinositol-anchors [1]. The complex GDNF-GFRα1 associates to a single transmembrane RET receptor tyrosine kinase, promoting dimerization and activation of RET’s intracellular kinase domain, leading to stimulation of multiple downstream pathways [1]. In the mammalian testes, GDNF is produced by testicular somatic cells, including Sertoli cells [2, 10, 11], peritubular myoid cells under influence of androgens [12, 13] and testicular endothelial cells which seem the major GDNF-producing source in the mouse testis [14]. GFRα1 is expressed in a subpopulation of single type A spermatogonia (A_s_) (inhibitor of DNA binding 4 +) which is considered the purest functional SSC population in mice [15, 16]. GFRα1 is not exclusively expressed in SSCs, but it is also detected in types A paired (A_pr_) and aligned (A_al_) spermatogonia in the mouse testis [17–21]. Similar expression pattern for GFRα1 has been reported in the testes of numerous mammalian species, such as hamster [22], pig [23], collared peccary [24, 25], buffalo [26], different equine species [27], and primates including humans [28, 29]. The mechanisms underlying the regulation of GDNF, especially in Sertoli cells, are not fully understood. This lack of knowledge is partially attributed to difficulties on working with adult primary Sertoli cells and due to the absence of efficient *ex vivo* organ culture systems that conserve adult Sertoli cell functions (see review in Parekh et al [8] and Mäkelä and Hobbs [9].

In nonmammalian species, particularly in the group of fish, Gdnf/Gfrα1 homologs have been found in a limited number of species, such as dogfish (*Scyliorhinus canicula*) [30, 31], rainbow trout (*Oncorhynchus mykiss*) [32–34] and medaka (*Oryzias latipes*) [35]. Unlike mammals, Gdnf and Gfrα1 were found co-expressed in type A undifferentiated spermatogonia of the above-mentioned species, suggesting an autocrine mechanism for Gdnf-mediated functions in fish testes [32]. The physiological relevance of Gdnf for type A undifferentiated spermatogonia has been further demonstrated by *in vitro* studies showing that recombinant human GDNF promoted proliferation and long-term maintenance of dogfish spermatogonia with stem characteristics [31]. Similar findings were found by Wei et al. [36] who showed that two Gdnf homologs in medaka, named Gdnfa and Gdnfb, stimulated proliferation of SG3, a spermatogonial stem cell line derived from adult medaka. On the other hand, studies in rainbow trout revealed that *gdnfb* mRNA levels increased during the arrest of spermatogenic cycle (end of germ cell proliferation and differentiation), suggesting that Gdnfb is likely involved in the repression of SSC differentiation rather than proliferation [33]. Therefore, more studies are needed to unravel the possible autocrine/paracrine roles of Gdnf on SSC niche in fish, aiming to expand our knowledge about the critical factors involved in the SSC activity in these animals, as well as predicting the consequences of changes involved in the physiological mechanisms related to the Gdnf. According to Oatley and Brinster [37], the reduction or loss of SSC function disrupts spermatogenesis leading to subfertility or infertility in males and, therefore, knowing the mechanisms that regulate SSC homeostasis is imperative for the conservation of species or for its use as an experimental model in studies focusing on the treatment of pathological conditions of the reproductive organs of humans.

Considering this background and the lack of knowledge on Gdnf-Gfrα1 signaling in the zebrafish testis, this study initially performed phylogenetic and conserved synteny analysis of Gfrα1 followed by expression profiling of *gdnf* (*gdnfa* and *gdnfb*) and *gfrα1* (*gfrα1a* and *gfrα1b*) transcripts in the zebrafish testes. Subsequently, the cellular types expressing Gdnf and Gfrα1 were identified in the zebrafish testis, and the biological effects of Gdnf were assessed using an *ex vivo* testis culture system established for zebrafish. To the best of our knowledge, the present study was pioneer in evidencing that Gdnf is expressed in the germ cells of zebrafish, whereas its co-receptor, Gfrα1, was detected in Sertoli cells and among different types of spermatogonia, in which signal was more intense in type A undifferentiated spermatogonia. Moreover, we showed that Gdnf, a germinal signal, exerts autocrine and paracrine roles in the regulation of zebrafish testicular function through stimulating survival of type A spermatogonia, and inducing mitosis of Sertoli cells, respectively.

## 2 Material and methods

### 2.1 Zebrafish stocks

Sexually mature zebrafish (*Danio rerio*, outbred) (4-5 months old) were bred and raised in the aquarium facility of the Department of Structural and Functional Biology, Institute of Biosciences, São Paulo State University (Botucatu, Brazil). Fish were kept in tanks of 6-L in the recirculating system and temperature conditions similar to the natural environment (27°C) under proper photothermal conditions (14 hours of light and 10 hours dark). Salinity, pH, dissolved oxygen and ammonia were monitored in all tanks every day. Fish were fed twice a day using commercial food (Zeigler®, Gardners, PA, USA). No mortality was observed during experiments. Handling and experimentation were consistent with Brazilian legislation regulated by National Council for the Control of Animal Experimental (CONCEA) and Ethical Principles in Animal Research of São Paulo State University (Protocol n. 666-CEUA). Zebrafish is a tropical freshwater fish natural to rivers in Southern Asia, mainly in Northern India, Pakistan, Bhutan and Nepal [38–40], has been considered a versatile model for reproductive biology [41], besides being used as a model for translational research on human health and disease [42]. Therefore, these aspects justify the choice of this species in our study.

### 2.2 Sequence analysis

The predicted amino acid sequences for Gfrα1a and Gfrα1b of *D. rerio* (Q98TT9 and Q98TT8, respectively), GFRA1 of *Homo sapiens* (P56159), *Rattus norvegicus* (Q62997) and *Mus musculus* (P97785) were obtained from The Universal Protein Resource (UniProt, accessed 09/12/2019), and aligned using the MEGA algorithm allocated on the Geneious Pro 4.8.5 software [43]. For the phylogenetic analysis, we retrieved the protein sequences of GFRα1 (Gfrα1a and Gfrα1b) from The Universal Protein Resource (UniProt, accessed 02/25/2020), from National Center for Biotechnology Information (NCBI, 02/25/2020) and Ensembl (accessed 02/25/2020 [44]). For this analysis, we retrieved vertebrate sequences for GFRα1 and Growth arrest-specific protein 1 (GAS1) from human (GAS1 as an outgroup) (**Figure 2**). The predicted amino-acid sequences were aligned using the Muscle algorithm [45] allocated on the Geneious Pro 4.8.5 software [43]. The choice of the best-fit model of evolution was performed with SMS [46]. Phylogenetic reconstruction was determined by Bayesian methods implemented in Beast v1.7.0 software [47]. This step was carried out according to Geraldo et al. [48], with adaptations. Brunch values were supported by posterior probabilities obtained by Bayesian analysis. For Bayesian method generations, the burn-in was determined in Tracer [47] through log likelihood scores, and data were summarized in TreeAnnotator [47] after trees that were out of the convergence area had been discarded. The visualization and the final tree edition were performed using FigTree v1.3.1 [47]. In the phylogenetic analyses, the proportion of invariable sites and γ-distributed rate variation across sites were estimated, and the substitution of rate categories were set in four categories. The parameters set to reconstruct the phylogeny are shown in **Table S2**. To construct the syntenic regions of *GFRA1* (human), *Gfr*α*1* (rat and mouse), *gfrα1a* and *gfrα1b* (zebrafish) genes, we used the GenBank database, available at the National Center for Biotechnology Information (NCBI; http://www.ncbi.nlm.nih.gov/) and Ensembl [44].

### 2.3 Expression profiling of *gdnf* (*gdnfa* and *gdnfb*) and *gfrα1* (*gfrα1a* and *gfrα1b*) transcripts in the zebrafish testes

To investigate the transcript abundance of *gdnf* (*gdnfa* and *gdnfb*) and *gfrα1* (*gfrα1a* and *gfrα1b*) in the zebrafish testes, total RNA from testes (n = 4 males) was extracted by using the commercial RNAqueous®-Micro kit (Ambion, Austin, USA), according to manufacturer’s instructions. The cDNA synthesis was performed as usual procedures [49]. Quantitative reverse transcription polymerase chain reaction (RT-qPCR) was conducted using 10 µL 2× SYBR-Green Universal Master Mix, 2 µL of forward primer (9 mM), 2 µL of reverse primer (9 mM), 1 µL of DEPC water, and 5 µL of cDNA. The relative mRNA levels of *gdnfa* (glial cell-derived neurotrophic factor a), *gdnfb* (glial cell-derived neurotrophic factor b), *gfrα1a* (gdnf family receptor alpha 1a) and *gfrα1b* (gdnf family receptor alpha 1b) (Cts) were normalized by the reference genes *ef1*α (elongation factor 1 α) and β*-actin*, expressed as relative values of control group (as fold induction), according to the 2^−(ΔΔCT)^ method. Primers were designed based on zebrafish sequences available at Genbank (NCBI, https://www.ncbi.nlm.nih.gov/genbank/) (**Table 1**).

**Table 1.**
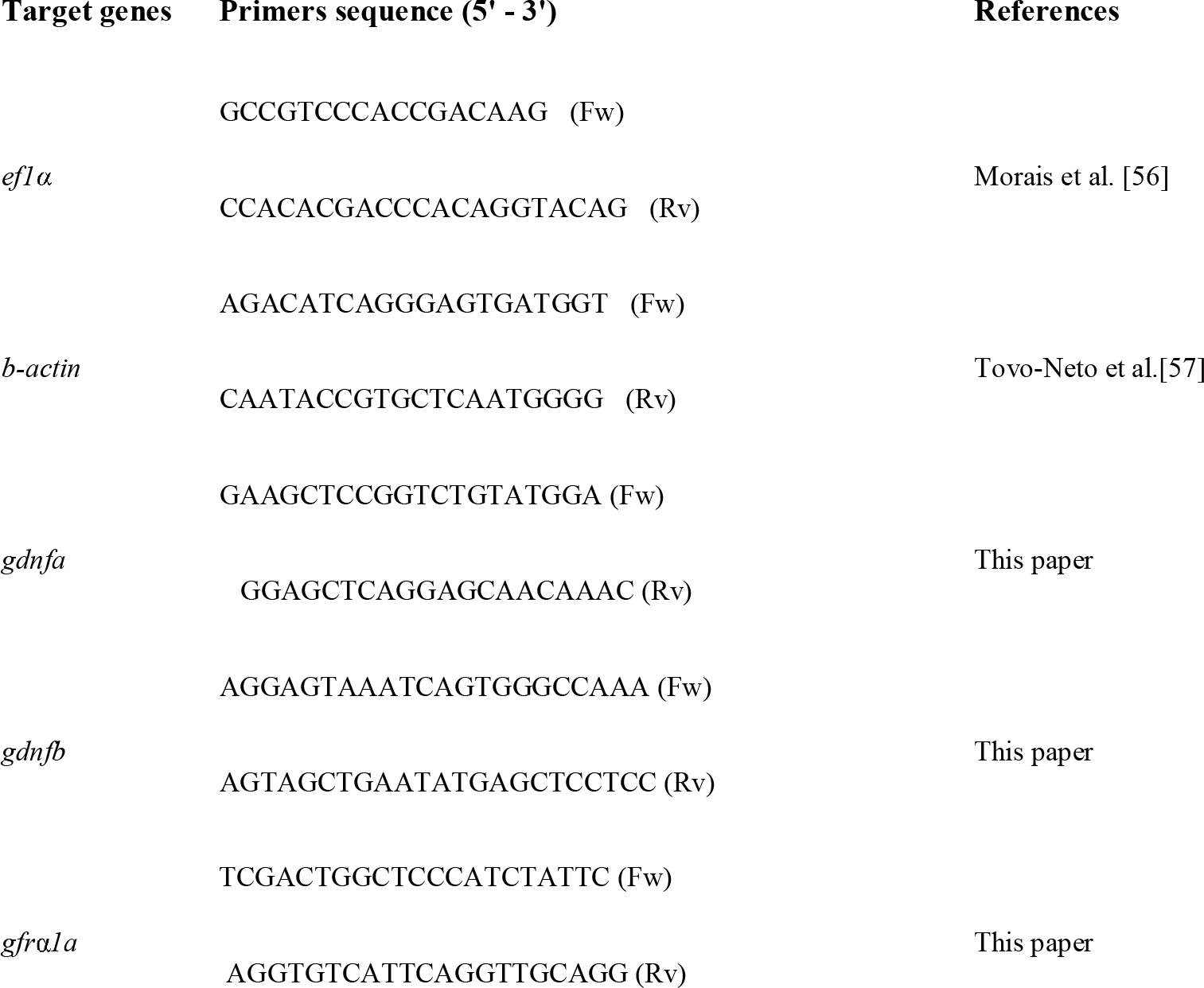
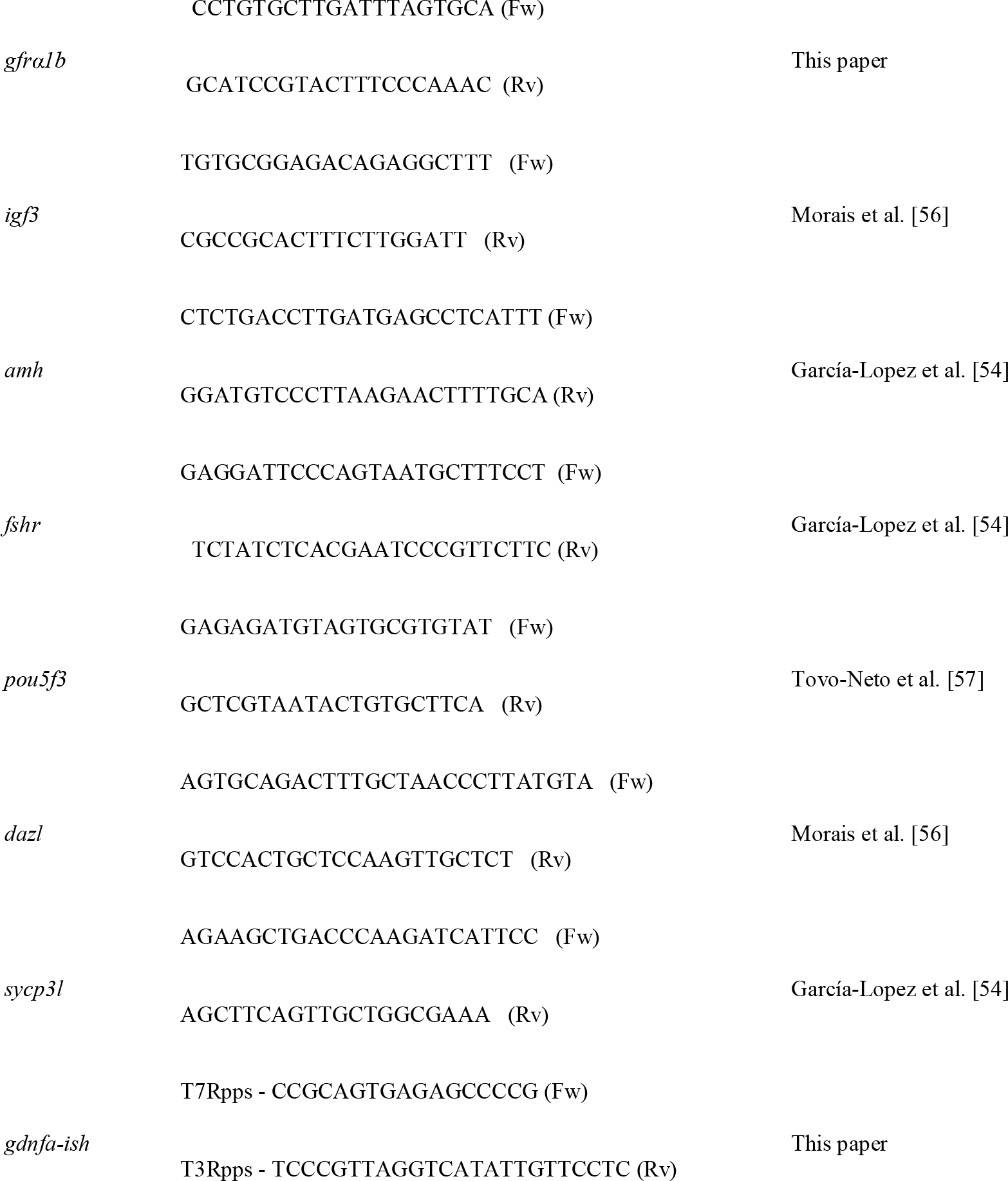
Primers used for gene expression analysis (RT-qPCR) and to generate DNA templates for digoxigenin (DIG)-labeled cRNA probe synthesis for *in situ* hybridization (ISH) (**Supplemental material**). Fw, forward; Rv, reverse; T7Rpps – T7 RNA polymerase promoter sequence at its 5’-end (5’ CCGGGGGGTGTAATACGACTCACTATAGGG-3’), T3Rpps – T3 RNA polymerase promoter sequence at its 5’-end (T3’GGGCGGGTGTTATTAACCCTCACTAAAGGG-3’).

### 2.4 Differential plating method

To identify the testicular cell fractions (germ or somatic cell enriched fractions) expressing *gdnfa* in zebrafish, a differential plating method was carried out as previously described by Hinfray et al. [50]. To this end, testes (n = 20 males) were digested with 0,2% collagenase (Sigma Aldrich, San Luis, MI, USA) and 0,12% dispase (Sigma Aldrich, San Luis, MI, USA), as described previously [49]. The total cell suspension was then submitted to a differential plating method, where the somatic cells adhere to the bottom of the plate, while germ cells either remain in suspension after 2–3 days of culture or only weakly associated with the firmly adhering somatic cells [50]. By using this method, germ and somatic cell enriched fractions can be obtained [50]. Total RNA from the cell suspensions (total, germ cell enriched, and somatic cell enriched) was extracted using PureLink® RNA Mini Kit (Ambion, Austin, TX, USA), according to manufacturer’s instructions. After cDNA synthesis using SuperScript® II Reverse Transcriptase kit (Invitrogen, Carlsbad, CA, USA) and random hexamers, the relative mRNA levels of *pou5f3* (POU domain, class 5, transcription factor 3) (spermatogonia marker) and *gdnfa* were determined by qRT-PCR. β*-actin* and *ef1* were used as housekeeping genes. The quantification cycle (Cq) values were determined in a StepOne system (Life Technologies, Carlsbad, CA, USA) using SYBR Green (Invitrogen, Carlsbad, CA, USA) and specific primers (**Table 1**). All RT-qPCR reactions (10-20 µL) used 900 nM for each primer (forward and reverse) and 300 ng of total cDNA. Each reaction was performed in duplicate and relative gene expression levels were calculated according to the 2^−(ΔΔCT)^ method.

### 2.5 Immunofluorescence and Western blot

Testes (n = 10 males) were fixed with 4% paraformaldehyde in PBS (Phosphate Buffered Saline) (1X, pH 7.4) for 1 hour, embedded in paraplast (Sigma Aldrich, San Luis, MI, USA) and sectioned at 5 μm thickness. After deparaffinization and rehydration, sections were submitted to antigen retrieval by heating slides in sodium citrate buffer (10 mM sodium citrate, 0,05% Tween 20, pH 6.0) until temperature reaches 95-100°C in a microwave. To reduce background fluorescence, slides were incubated with NaBH4 (sodium borohydride - 0.01g dissolved in 1 mL of distilled water) (Sigma Aldrich, San Luis, MI, USA) for 3 minutes. Subsequently, slides were rinsed with 1X PBS (pH 7.4) and incubated with the biotinylated primary antibody rabbit anti-zebrafish Gfrα1a (1:300, 1X PBS pH 7.4) at 4°C overnight. Zebrafish polyclonal biotinylated antibody anti-Gfrα1a was synthesized by Rheabiotech (Campinas, Brazil) using the specific antigen sequence ‘RLDCVKANELCLKEPGCSSK’ located at the N-terminus of zebrafish Gfrα1a (**Figure 1**). This antibody is also potentially able to recognize other Gfrα1 isoforms, such as GFRA1 of humans, rodents, and Gfrα1b of zebrafish (**Figure 1**). After rising, the slides were incubated with Dylight 488 Streptavidin (BioLegend®-San Diego, CA, USA) (1:400) or Alexa Fluor 594 Streptavidin (BioLegend®-San Diego, CA, USA) (1:400) in 1X PBS (pH 7.4) for 60 minutes at room temperature. Subsequently, sections were counterstained with Hoechst (1:2000, 1X PBS pH 7.4) (Invitrogen, Carlsbad, CA, USA), or Propidium iodide (PI) (BioLegend®-San Diego, CA, USA) (1 mg PI to 1 mL distilled water), and mounted with ProLong Gold Antifade (Thermo Fisher Scientific, Waltham, MA, USA). Control sections were prepared by preadsorbing the zebrafish Gfrα1a antibody with the corresponding peptide (10 μg/1 µL of antibody, Rheabiotech, Campinas, Brazil) or by omitting the primary antibody. Slides were photographed using a Leica SP5 laser scanning confocal microscope (Leica, Wetzlar, Germany) from the Electron Microscopy Center, Institute of Biosciences, São Paulo State University (Botucatu, Brazil) and germ cells were classified based on morphological criteria established by Leal et al. [51].

**Figure 1.**
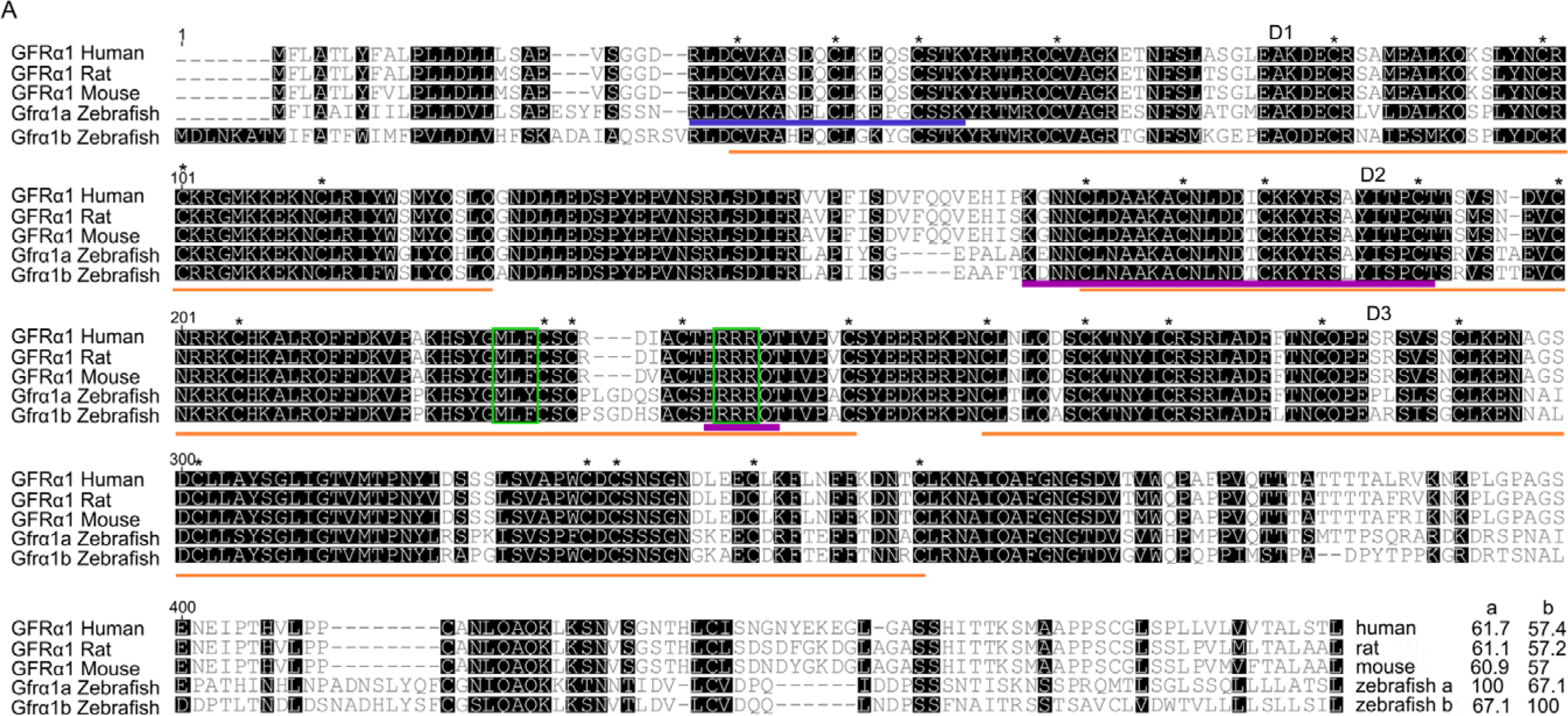
GFRα1 predicted amino acid sequence alignment. Numbers on the left top indicate amino acid positions, dashes indicate deletions, and black boxes indicate shared sequences. The three cysteine-rich domains (D1-3), 28 cysteine residues (*) (plus 2 in the terminal region) and two triplets (MLF and RRR) (green boxes) are highly conserved among human, rodents and zebrafish. At the end of the alignment are the percentage identity values of zebrafish Gfrα1a and Gfrα1b to the other corresponding sequences. The blue box indicates the amino acid sequence recognized by anti-zebrafish Gfrα1a antibody (Rheabiotech, Campinas, Brazil - see Material and Methods section). Purple line indicates the putative motifs critical for GFRα1 binding to GDNF and eliciting downstream signal transduction.

For the Western blot analysis, testes (n = 10 males) were homogenized in an extraction TBST buffer (10 mM Tris–HCl, pH 7.5; 150 mM NaCl; 0,1% Tween 20) containing a cocktail of protease inhibitors (Roche Applied Science, Mannheim, Germany). Subsequently, the homogenate was incubated on ice for 15–20 minutes before sonication (3× 1 minute on ice), and centrifuged at 4000 rpm at 4°C for 20 minutes in order to determine the total protein concentration through NanoVue spectrophotometer (GE Healthcare, Chicago, IL, USA). A total of 40 µg protein was analyzed by sodium dodecyl sulfate polyacrylamide gel electrophoresis (SDS-PAGE). Protein extracts were blotted onto a nitrocellulose membrane (Amersham, Little Chalfont, UK), blocked with 3% non-fat milk diluted in 1X Tris-buffered saline (TBS) (150 mM NaCl, 50 mM Tris-HCl, pH 7.6.) for 1 hour, and incubated with the primary antibody rabbit anti-zebrafish Gfrα1a (1:500, Rheabiotech, Campinas, Brazil) at 4°C overnight. The membrane was washed with TBS and incubated with horseradish peroxidase-conjugated anti-rabbit IgG (1:5000, Santa Cruz Biotechnologies, Dallas, TX, EUA) for 2 hours. After washing, blots were developed with chemiluminescence substrate kit (Pierce ECL Western Blotting Substrate-GE Healthcare, Chicago, IL, USA) and the signal was captured by a CCD camera (ImageQuant LAS 4000 mini®, GE Healthcare, Chicago, IL, USA). As controls, some membranes were alternatively incubated with a primary antibody that has been preadsorbed with the respective peptide, as described above.

### 2.6 Recombinant human GDNF

To evaluate the effects of Gdnf on zebrafish spermatogenesis (see below), a recombinant human GDNF (rhGDNF) purchased from PeproTech® (London, UK) (catalog number 450-10; https://www.peprotech.com/en/recombinant-human-gdnf#productreviews) was used. The recombinant hormone was dissolved in sterile Lebovitz medium (L-15) (Sigma-Aldrich, St. Louis, USA) at the concentration of 100 µg/mL, and subsequently aliquoted and stored at -20° C until use. After identifying the binding sites between rhGDNF and human GFRA1A, a 3D structure model was built to predict the interaction sites between rhGDNF and zebrafish Gfrα1a (Q98TT9). The 3D protein structure used was obtained through SWISS-MODEL (swissmodel.expasy.org) with multiple target sequences representing different subunits of a hetero-oligomer (hetero-2-2-mer), and analyzed the quality of the modeling with the Ramachandran allocated in Rampage software [52]. The template (4ux8.1) and the final model were viewed in the software Pymol (The PyMOL Molecular Graphics System, Version 1.8 Schrödinger, LLC).

### 2.7 Testis tissue culture

The effects of rhGDNF on zebrafish spermatogenesis were investigated using a previously established *ex vivo* culture system [53]. After dissecting out the testes (paired structure) (n = 30 males), each testis (left and right) was placed on a nitrocellulose membrane measuring 0.25 cm^2^ (25 µm of thickness and 0.22 µm of porosity) on top of a cylinder of agarose (1.5% w/v, Ringer’s solution—pH 7.4) with 1 mL of culture medium into a 24-well plate. In this system, one testis (left) was incubated in the presence of rhGDNF (100 ng/mL - based on Gautier et al [55]) and its contra-lateral one (right) in a basal culture medium (L-15). The medium was changed every 3 days of culture. After 7 days, testes were collected for histomorphometrical analysis, BrdU (bromodeoxyuridine) incorporation assay, and gene expression (RT-qPCR) (see below). Additional cultures were carried out to assess the interaction of Gdnf with Fsh-mediated effects on zebrafish spermatogonial phase [49]. To this end, zebrafish testes (n = 10 males) were incubated with recombinant zebrafish Fsh (rzfFsh) (100 ng/mL [53]) (U-Protein Express B.V; Utrecht, the Netherlands) in the presence or absence of rhGDNF (100 ng/mL) for 7 days. After the culture period, testes were collected for gene expression analyses (RT-qPCR).

For histomorphometrical analysis, zebrafish testicular explants (n =10) were fixed in 4% buffered glutaraldehyde at 4 °C overnight, dehydrated, embedded in Technovit (7100-Heraeus Kulzer, Wehrheim, Germany), sectioned at 4µm thickness, and stained with 0.1% toluidine blue to quantify the different germ cell types at 40× and 100× objectives using a high-resolution light microscope (Leica DM6000 BD, Leica Microsystems, Wetzlar, Germany). In this analysis, five histological fields for each animal were randomly selected for counting the frequency of germ cell cysts [type A undifferentiated spermatogonia (A_und_), type A differentiated spermatogonia (A_diff_), type B spermatogonia (SPG B), spermatocytes (SPC), and spermatids (SPT)], as previously described [49, 56].

To evaluate the effects of rhGDNF on germ cell proliferation, 100 µg/mL BrdU (Sigma Aldrich, San Luis, MI, USA) was added to the culture medium during the last 6 hours of incubation. After incubation, zebrafish testes (n = 10) were fixed at 4 °C overnight in freshly prepared methacarn (60% [v/v] absolute ethanol, 30% chloroform, and 10% acetic acid) for 4 h. Subsequently, testes were dehydrated, embedded in Technovit 7100 (7100-Heraeus Kulzer, Wehrheim, Germany), sectioned at 4µm thickness, and submitted to BrdU immunodetection, as previously described [51, 56]. The mitotic index or BrdU incorporation ratio of types A_und_, A_diff_ and Sertoli cells was performed as described previously [51,56,57].

For the gene expression analyses, total RNA from testicular explants (n = 20 males) was extracted by using the commercial RNAqueous®-Micro kit (Ambion, Austin, USA), according to manufacturer’s instructions. The cDNA synthesis was performed as described above. RT-qPCR reactions were conducted using 10 µL 2× SYBR-Green Universal Master Mix (Bio-Rad, Hercules, CA, USA), 2 µL of forward primer (9 mM), 2 µL of reverse primer (9 mM), 1 µL of DEPC water, and 5 µL of cDNA. The relative mRNA levels of *gdnfa* (glial cell-derived neurotrophic factor a), *gfrα1a* (gdnf family receptor alpha 1a), *gfrα1b* (gdnf family receptor alpha 1b), *amh* (anti-Müllerian hormone), *igf3* (insulin-like growth factor 3), *fshr* (follicle stimulating hormone receptor), *pou5f3* (POU domain, class 5, transcription factor 3), *dazl* (deleted in azoospermia-like) and *sycp3l* (synaptonemal complex protein 3) were evaluated. The mRNA levels of the targets (Cts) were normalized by the reference gene β-actin, expressed as relative values of basal expression levels, according to 2^−(ΔΔCT)^ method. Primers were designed based on zebrafish sequences available at Genbank (NCBI, https://www.ncbi.nlm.nih.gov/genbank/) (**Table 1**).

### 2.8 *In silico* analysis of putative regulatory sequences upstream human *GDNF*, mouse *Gdnf* and zebrafish *gdnfa*

To retrieved the putative regulatory sequences upstream human *GDNF* (NM_000514.4), mouse *Gdnf* (NM_010275.3) and zebrafish *gdnfa* (NM_131732.2), the transcription start site (TSS) was found in the Eukaryotic Promoter Database (EPD) and the promoter regulatory regions (3’to 5’) was prospected by the flanking regions (2000 bp) extracted from National Center for Biotechnology Information (NCBI). The cAMP response elements (CRE, 4 different sequences), androgen receptor binding site (AR, full and half sequences), several NF-kB-binding sites, N-Box, E-Box, TATA-Box, and GC-Box (**Table S3**) were prospected using sequences described in the literature [8, 59–64].

### 2.9 Statistical analyses

Graphpad Prism 7.0 (Graphpad Software, Inc., San Diego, CA, USA, http://www.graphpad.com) was used for all statistical analysis. Data were initially checked for deviations from variance normality and homogeneity, before the analysis. Data normality was assessed through Shapiro-Wilks test, and variance homogeneity was assessed through Bartlette’s test. Significant differences between two groups were identified using paired Student’s t-test, at 5% probability. Comparisons of more than two groups were performed with one-way ANOVA followed by Student-Newman-Keuls test, at 5% probability.

## 3. Results

### 3.1 Sequence analyses, phylogenetic tree and genomic organization of zebrafish Gfrα1a and Gfrα1b

Sequence analysis revealed that both the predicted zebrafish Gfrα1a and Gfrα1b have sequence characteristics of Gfrα family members, such as the three cysteine-rich domains (D1-3), 28 cysteine residues (plus 2 in the terminal region), and two triplets (MLF and RRR) in the domain D2 (**Figure 1**). Sequence alignment of zebrafish Gfrα1a and 1b with different GFRA1s (human and rodents) revealed that the three cysteine-rich domains (D1, D2, D3) are highly conserved among the species, highlighting, in particular, the conserved residues and motifs in the domain D2 critical for GFRα1 binding to GDNF and eliciting downstream signal transduction as shown in mammals (**Figure 1**). Sequence analyses also demonstrated that zebrafish Gfrα1a and 1b have a 67,1% identity to each other, and the Gfrα1a showed higher identity with mammalian GFRA1 (61,7%, 61,1% and 60,9% similarity to human, rat and mouse GFRA1, respectively) when compared to Gfrα1b (57,4%, 57,2% and 57% identity to human, rat and mouse GFRA1, respectively) (**Figure 1**).

Phylogenetic analysis further confirmed that both zebrafish Gfrα1a and Gfrα1b are related to other fish Gfrα1a and Gfrα1b predicted sequences, respectively, and these isoforms diverge and form two separate fish-specific subclades (estimated posterior probability = 1) (**Figure 2A**). On the other hand, the GFRA1 sequences from other vertebrates (mammals, birds, reptiles, amphibians and Chondrichthyes) are clustered and form a separate clade from the fish Gfrα1 (estimated posterior probability = 0.851) (**Figure 2A**).

**Figure 2.**
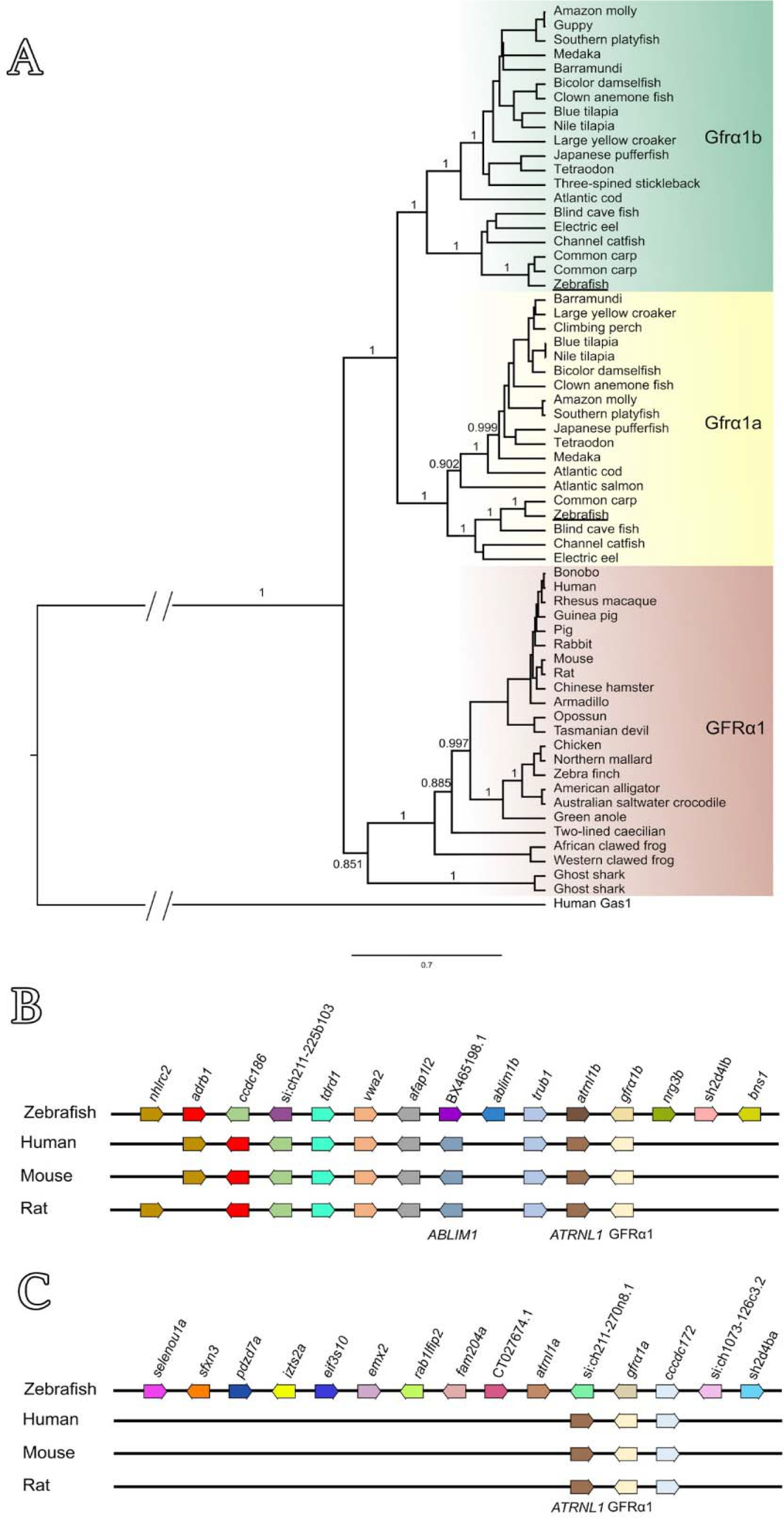
(A) Phylogenetic analysis of GFRα1 predicted amino acid sequences across vertebrates. Zebrafish Gfrα1a and 1b (both underlined) are clustered with other fish-specific Gfrα1a (yellow box) and Gfrα1b (green box) sequences, respectively, forming two separate subclades. Note that the GFRA1 sequences from other vertebrates (mammals, birds, reptiles, amphibians and Chondrichthyes) formed a separate clade (brown box). Branch values represent posterior probabilities obtained by Bayesian analysis (see **Table S1**). (B-C) Genomic organization and synteny comparisons among human *GFRA1*, rodents *Gfrα1* and zebrafish *gfrα1b* (B) or zebrafish *gfrα1a* (C). The syntenic regions were analyzed based on the alignment of the target genes and genomic annotation according to the GenBank database, available at National Center for Biotechnology Information and Ensembl.

A cross-species comparison of chromosome neighboring genes revealed that both of zebrafish *gfrα1a*- and *gfrα1b*-containing regions are syntenic to human *GFR*A*1*- and rodents *Gfr*α*1*- containing regions (**Figure 2B**). This analysis also showed that zebrafish *gfrα1b* gene (chromosome 12, NC_007123.7) showed a larger group of syntenic genes (8 out of 14 genes analyzed) when compared to zebrafish *gfrα1a* (chromosome 13, NC_007124.7) (2 out of 14 genes analyzed) (**Figure 2B**).

### 3.2 Expression profiling of *gdnf* (*gdnfa* and *gdnfb*) and *gfrα1* (*gfrα1a* and *gfrα1b*) in the zebrafish testes and identification of *gdnfa expressing cells*

RT-qPCR analyses revealed that both *gfrα1a* and *gfrα1b* were expressed in the zebrafish testes, and the *gfrα1a* transcripts were significantly more expressed than *gfrα1b* (**Figure 3**). With regard to Gdnf ligands (Gdnfa and Gdnfb), transcript analysis showed that *gdnfa* was by far the most abundant ligand in the zebrafish testes as compared with *gdnfb* levels [Ct = 28,54 (Me) for *gdnfa* vs. Ct = 31,04 (Me) for *gdnfb*] (**Figure 3**).

**Figure 3.**
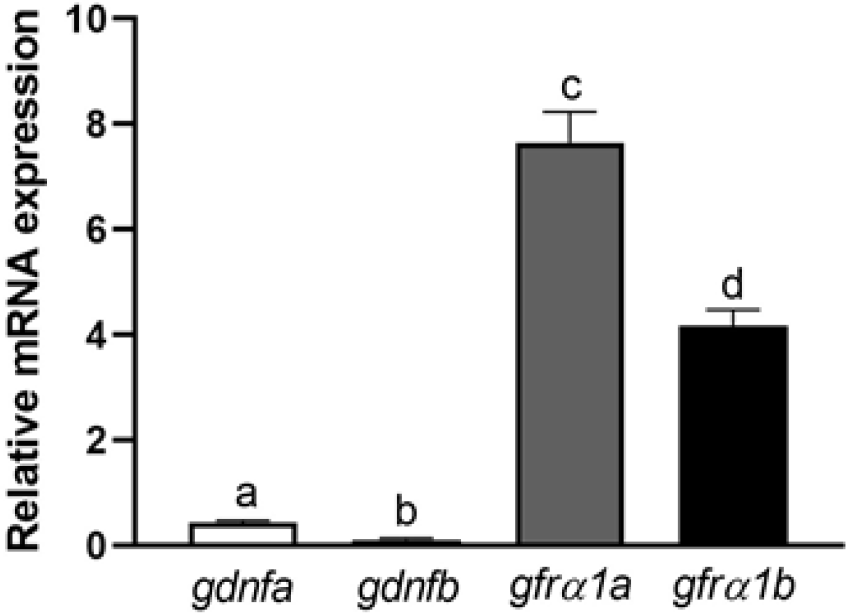
Relative mRNA expression levels of *gfdna*, *gdnfb*, *gfrα1a* and *gfrα1b* in the zebrafish testes. Bars represent the mean ± SEM (n = 4) and different letters denote significant differences among the evaluated genes (ANOVA followed by Student-Newman-Keuls test, *p* < 0.05).

Considering the transcript abundance of the Gdnf ligands, we attempted next to identify the cellular types expressing *gdnfa* mRNA in the zebrafish testes by employing *in situ* hybridization with specific antisense cRNA probe (**Table 1, Figure S1**), and RT-qPCR using RNA from isolated testicular cell populations (germ cell-enriched population and testicular somatic cells) (**Figure 4**). The first approach showed that *gdnfa* mRNA was expressed in germ cells at different stages of development (**Figure S1**). Nevertheless, due to limited resolution, it was not possible to unravel whether the signal was present or not in the Sertoli cells (**Figure S1**). This was attributed to the fact that cytoplasmic extensions of Sertoli cells protrude towards the lumen of a cyst in between the germ cells, making it difficult to accurately locate the signal. The precise identification of *gdnfa* expression sites was then accomplished through using testicular cell populations (germ cell-enriched population and testicular somatic cells) obtained after the differential plating method (**Figure 4A-C**). In this approach, RT-qPCR analysis showed an increase of *gdnfa* transcript levels in the germ cell-enriched population when compared to the levels found in the total testicular cell suspension (**Figure 4D**). Similar pattern of expression was found for *pou5f3*, a marker of types A_und_, A_diff_ and B spermatogonia (Souza, Doretto, Nóbrega - unpublished data), which confirmed the germ cell enrichment after the differential plating (**Figure 4D**). When analyzing the testicular somatic cell population, we found that *gdnfa* mRNA levels decreased significantly as compared to the levels observed in the germ cell-enriched fraction (**Figure 4E**).

**Figure 4.**
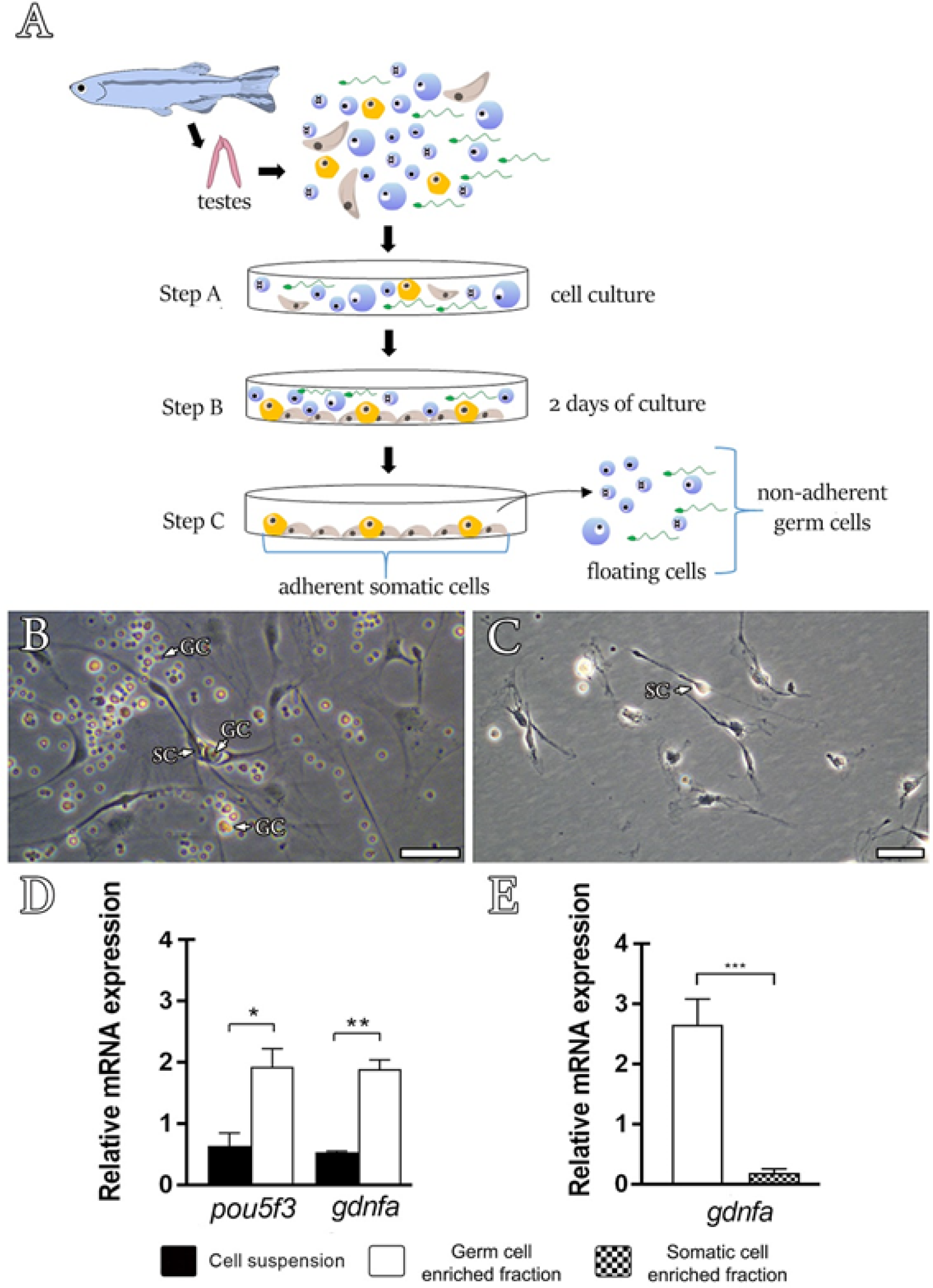
Differential plating method. (A) Scheme showing the steps of differential plating method according to Hinfray et al [50]. Briefly, a total testicular cell suspension obtained from enzymatic digestion was harvested (step A) in L-15 culture medium. After 2 days of culture, only somatic cells with adhesive properties (Sertoli cells, brown triangular shape; Leydig cells, yellow oval shape) adhere to the bottom of the plate (step B), while germ cells (non-adherent cells; blue shape cells) remain floating or loosely attached to the bottom of the plate (step C). After washing steps, it is possible to remove the germ cells (floating and also those germ cells weakly attached to the somatic cells), leaving the adherent somatic cells (Sertoli cells, Leydig cells, fibroblast and others) at the bottom of the plate. The firmly attached somatic cells can be obtained after extensive washing with trypsin. (B) Total testicular cell suspension after 2 days of culture. Note somatic adherent cells (SC) with cytoplasm extensions towards different germ cells (GC). (C) After extensive washing, somatic adherent cells (SC) remain attached to the bottom of the plate, while the floating and the weakly attached germ cells were removed. Scale bars: 20 µm. (D, E) Gene expression analysis of isolated zebrafish testicular cell populations: total cell suspension (black bar), germ cell enriched population (white bar), and testicular somatic cells (hatched bar). Cells were obtained from two independent experiments of differential plating method. Bars represent relative mRNA levels of target genes (*gdnfa* or *pou5f3*) expressed as mean ± SEM; asterisks indicate significant differences between the cell populations (unpaired t-test, \**p* < 0.05, \*\**p* < 0.01, \*\*\**p* < 0.001).

### 3.4 Localization of Gfrα1a protein in zebrafish testis

Immunofluorescence revealed that Gfrα1a protein was found in all generations of zebrafish spermatogonia, as types A undifferentiated (A_und_), differentiated (A_diff_) and B (SPG B) spermatogonia, but it was not detected in meiotic and post-meiotic germ cells, such as spermatocytes (SPC), spermatids (SPT) and spermatozoa (SPZ) (**Figure 5**). It is worthy to note that the staining pattern among the different generations of spermatogonia varied with the development stage (**Figure 5 A, C-E**). The Gfrα1a signal was finely dispersed in the cell surface and cytoplasm of type A_und_ spermatogonia (**Figure 5C**), and later, became more aggregated, forming intensely stained spots in type A_diff_ spermatogonia (**Figure 5D**). In type B spermatogonia, the Gfrα1a signal becomes finely dispersed again (Figure 5E), and gradually decreases as the number of spermatogonia B increases within the cyst, until being undetectable in the SPC cysts (**Figure 5A**). Furthermore, Gfrα1a was also found in Sertoli cells contacting germ cells in different stages of development (**Figure 5A, B**). The specificity of the antibody (anti-zebrafish Gfrα1a) was confirmed by immunoblots (**Figure 5F**), and control sections using either preadsorbed antibody with the corresponding peptide or omitting the antibody (**Figure S2**).

**Figure 5.**
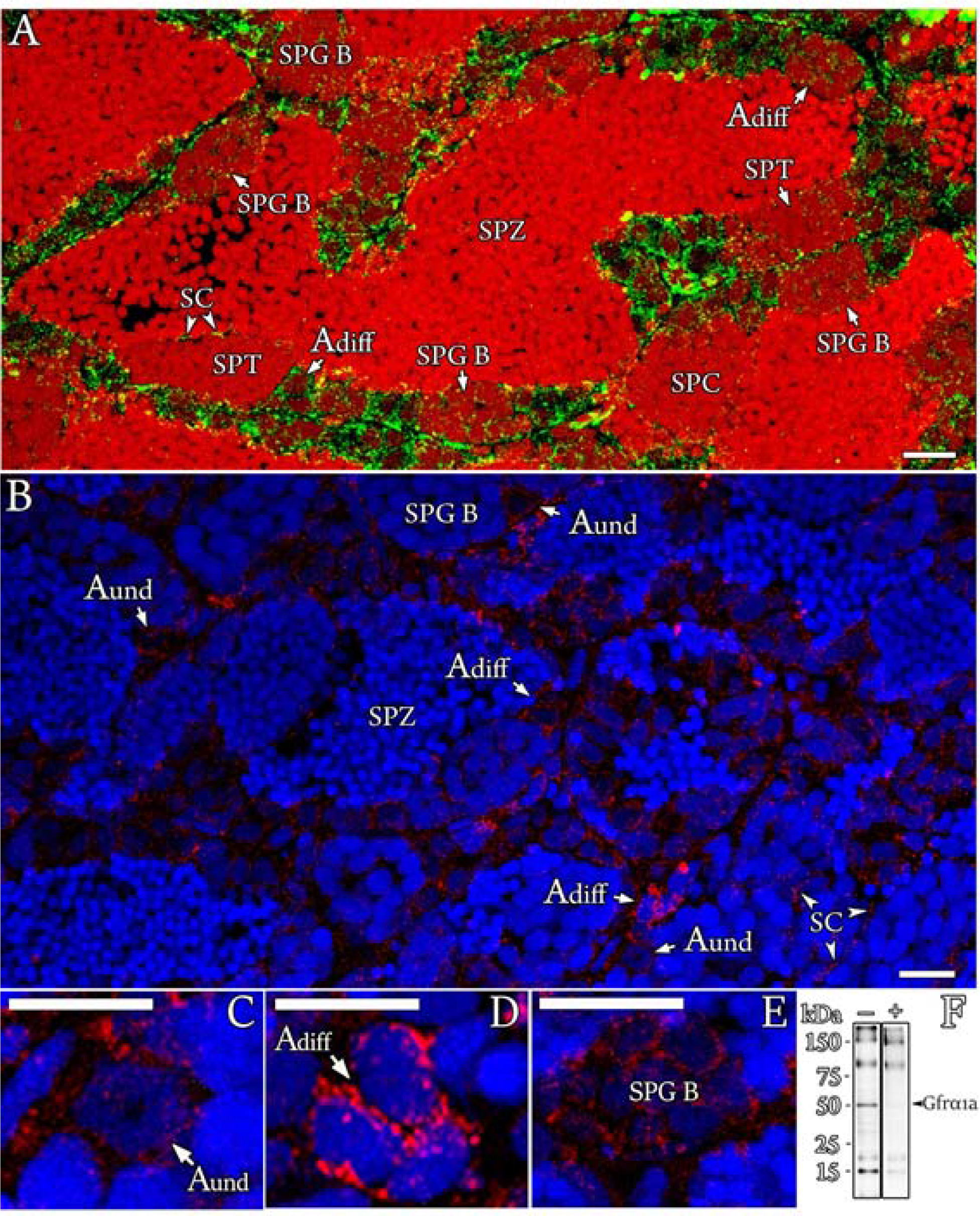
Cellular localization of Gfrα1a protein in zebrafish testis. (A-E) Immunofluorescence for Gfrα1a protein (green - A; red - B-E) on testis sections of sexually mature zebrafish. The spermatogonial generations, including type A undifferentiated spermatogonia (A_und_), type A differentiated spermatogonia (A_diff_) and type B spermatogonia (SPG B), were immunoreactive to Gfra1a, although staining pattern among them varied according the developmental stage. The signal was not found in spermatocytes (SPC), spermatids (SPT) and spermatozoa (SPZ). Note that Sertoli cells (SC) contacting germ cells in all stages of development were also immunoreactive to Gfrα1a. Cell nuclei were counterstained with propidium iodide (A) or Hoechst (B-E). Scale bars: 15 µm. (F) Gfrα1a (approximately 52 kDa - kilodaltons) immunoblots of whole testes with (+) or without (-) preadsorbed antibody, confirming the presence of the protein in the zebrafish testes and the antibody specificity.

### 3.5 3D model for predicting the interaction between rhGDNF and zebrafish Gfrα1a

To investigate the rhGDNF effects on zebrafish spermatogenesis, we first generated a 3D structure model to predict the possible interaction sites between human GDNF and zebrafish Gfrα1a (**Figure 6A, box2, box3**). The 3D structure (hetero-2-2-mer) was built according to the homology of the 4ux8.1 template, and showed a GMQE value of 0.63 with 74% of identity and a resolution of 24Å (method: Electron Microscopy) when compared to human GDNF-GFRA1 interaction (merged in the 3D structure) (**Figure 6A** **- box2, box3**). Moreover, the predictive model demonstrated that 89.8% of the amino acid residues were in the most favorable regions, 7% of residues were situated in allowed regions (∼2% expected), and 3.1% in the outlier regions according to Ramachandran plots. The 3D structures of the hetero-2-2-mer (GDNF-zebrafish Gfrα1a) were based on the homology modeling templates, and are shown in **Figure 6A** (**box2, box3**). A more detailed information regarding the predictive interaction model between GDNF and zebrafish Gfrα1a can be found at Supplementary Material (**Figure S3, Video S1**). Moreover, the alignment of zebrafish Gdnfa with rhGDNF showed conserved regions, particularly in the binding sites to human GFRA1 or zebrafish Gfrα1a (**Figure 6B**).

**Figure 6.**
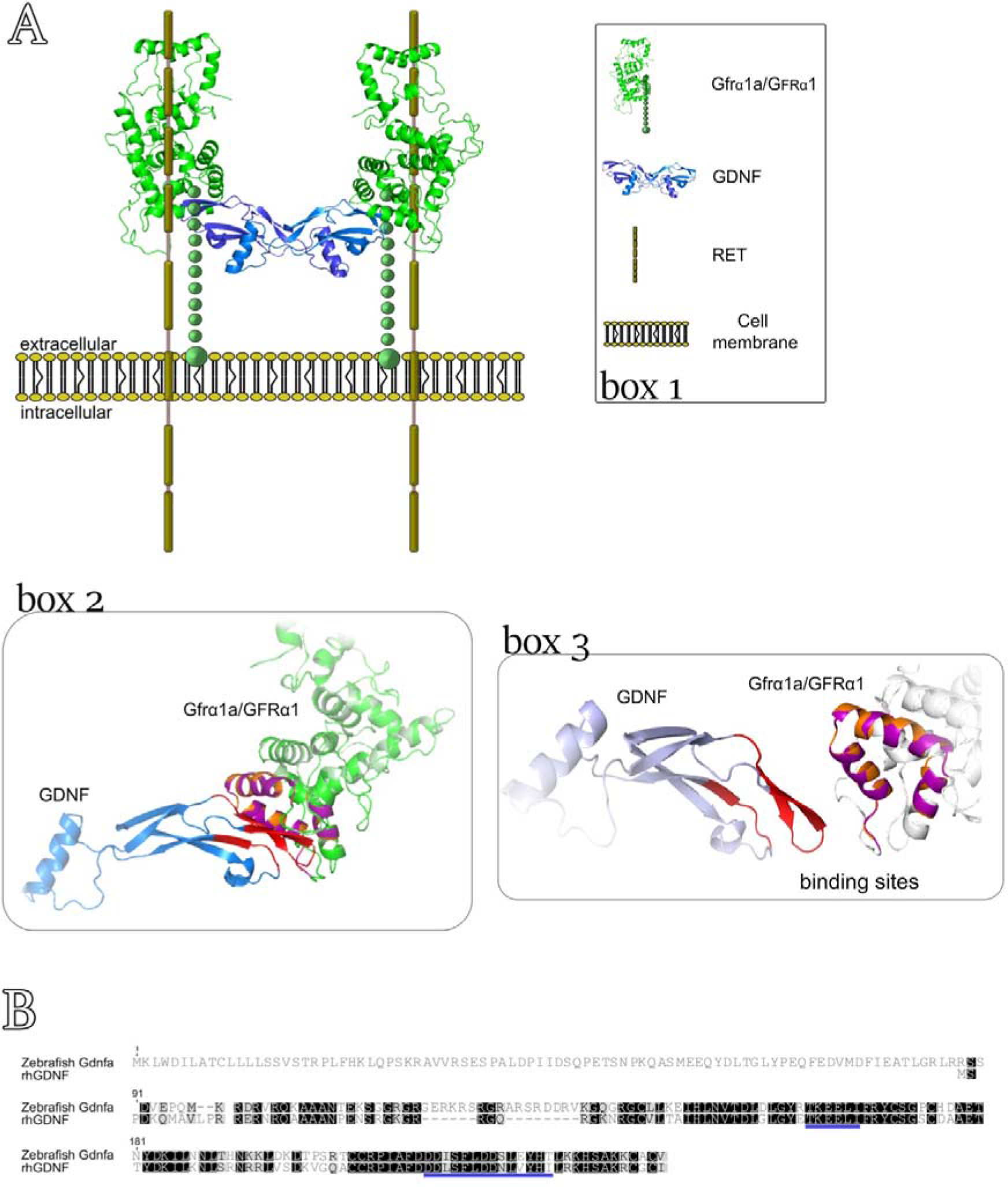
A 3D model to predict the interaction between rhGDNF and zebrafish Gfrα1a. (A) Box 1 depicts the molecular components of the complex GDNF-GFRα1-RET. Box 2 and 3 show the predictive 3D model (template 4ux8.1), in which the structural similarities between zebrafish Gfrα1a and human GFRA1 is represented by orange-purple, and the identity of the structure formed at the binding sites is indicated by red. In box 2, the color green shows the conserved amino-acid sequences between zebrafish Gfrα1a and human GFRA1, and blue indicates the GNDF protein. In box 3, we highlighted the interaction sites between human GDNF and zebrafish Gfrα1a/human GFRA1. (B) Alignment of zebrafish Gdnfa with rhGDNF. The blue lines indicate the conserved binding sites to zebrafish Gfrα1a or human GFRA1.

### 3.6 Biological effects of rhGDNF on spermatogenesis, cellular proliferation and gene expression analysis

To investigate the potential role of Gdnf in zebrafish spermatogenesis, we first examined whether rhGDNF could affect germ cell composition and cellular proliferation, using a previously established primary testis tissue culture system (**Figure 7A-D**). The results showed that rhGDNF (100 ng/mL) increased the abundance of types A_und_ and A_diff_ spermatogonia as compared to basal condition (**Figure 7C**). This data is also consistent with the proliferation activity of these cells showing that treatment with rhGDNF (100 ng/mL) augmented the mitotic index of both types of spermatogonia (A_und_ and A_diff_) as compared to their basal mitotic index (**Figure 7A, B, D**). Moreover, histomorphometrical analysis showed that rhGDNF decreased the frequency of type B spermatogonia, whereas no effects were seen for meiotic and post-meiotic germ cells (**Figure 7C**). In this study, we also quantified Sertoli cell proliferation (**Figure 7E**), reasoning that change in the proliferation of Sertoli cells associated with types A_und_ or A_diff_ spermatogonia would indicate creation of new niche space or supporting development of differentiating spermatogonia cysts, respectively [65]. Our results then demonstrated that treatment with rhGDNF stimulated Sertoli cell proliferation, particularly of those Sertoli cells associated with proliferating types A_und_ and A_diff_ spermatogonia (**Figure 7E**).

**Figure 7.**
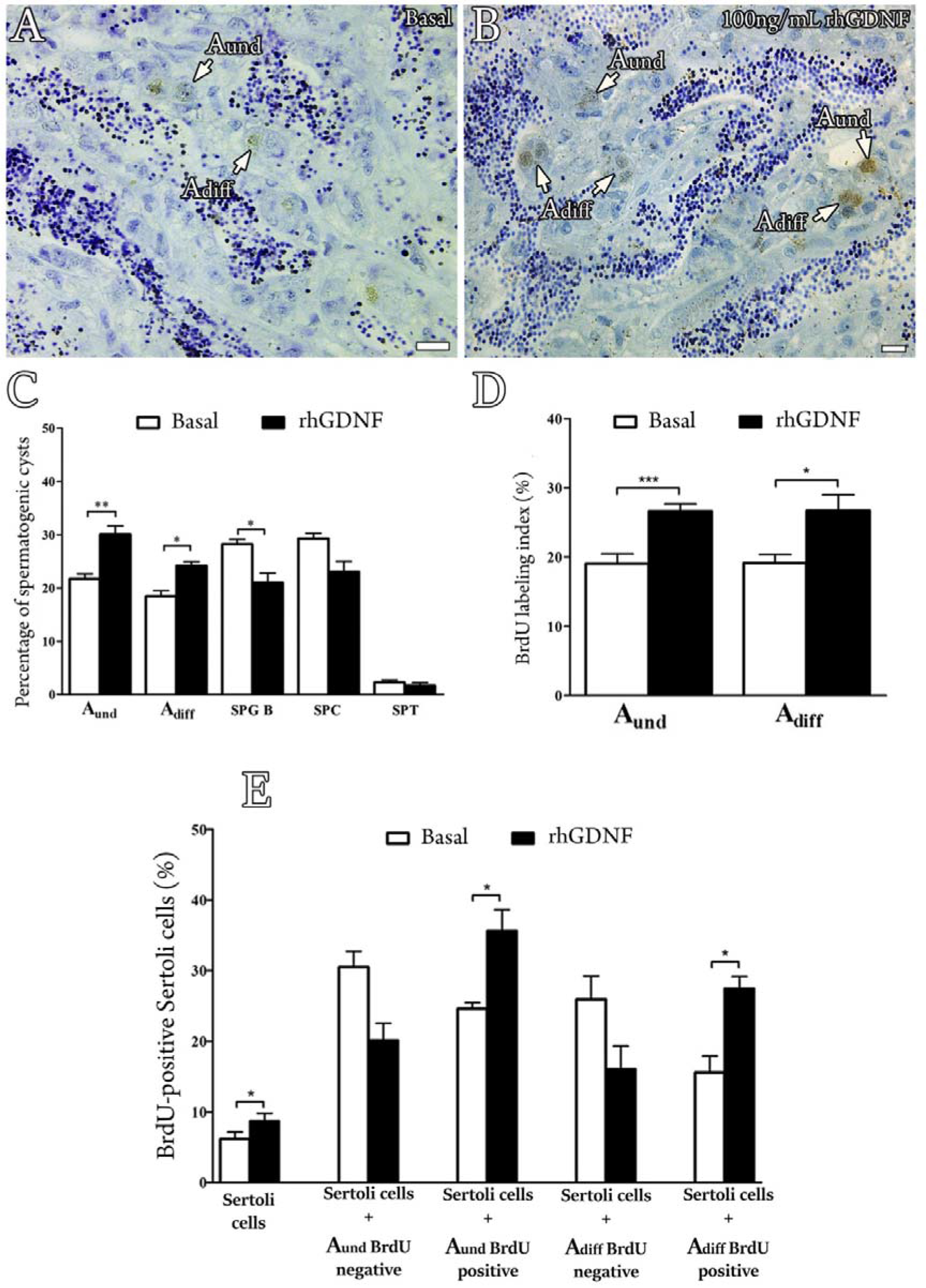
Effects of Gdnf on germ cell composition and cellular proliferation, using a previously established primary testis tissue culture system. (A-B) BrdU immunodetection from zebrafish testicular explants incubated for 7 days in the absence (Basal) or presence of rhGDNF (100 ng/mL), demonstrating a higher proliferation activity for type A undifferentiated spermatogonia (A_und_) and type A differentiated spermatogonia (A_diff_) in the presence of rhGDNF. (C) Cystic frequency of zebrafish testis explants after 7 days of incubation in the absence (Basal) or presence of rhGDNF (100 ng/mL). Types A_und_, A_diff_ and B spermatogonia (SPG B), spermatocytes (SPC) and spermatids (SPT) were counted. (D) Mitotic indices of type A_und_ and A_diff_ spermatogonia after incubation for 7 days in the absence (Basal) and presence of rhGDNF (100 ng/mL). (E) Mitotic indices of Sertoli cells in association with BrdU-negative or BrdU-positive type A_und_ and A_diff_ spermatogonia after incubation for 7 days in the absence (Basal) or presence of rhGDNF (100 ng/mL). Bars represent the mean ± SEM (n = 10). Paired t-test, *** *p* < 0.001; ** *p* < 0.01. Scale bars: 15 µm.

In order to elucidate the molecular mechanisms mediated by rhGDNF on basal or Fsh-induced spermatogenesis, we performed gene expression analyses of selected genes related with Gdnf signaling (*gdnfa*, *gfrα1a* and *gfrα1b*), Sertoli cell growth factors (*igf3* and *amh*), Fsh signaling (*fshr*) and germ cell markers (undifferentiated and differentiated spermatogonia - *pou5f3*; differentiated spermatogonia and preleptotene spermatocytes - *dazl;* and primary spermatocytes - *scyp3l)* (**Figure 8**).

**Figure 8.**
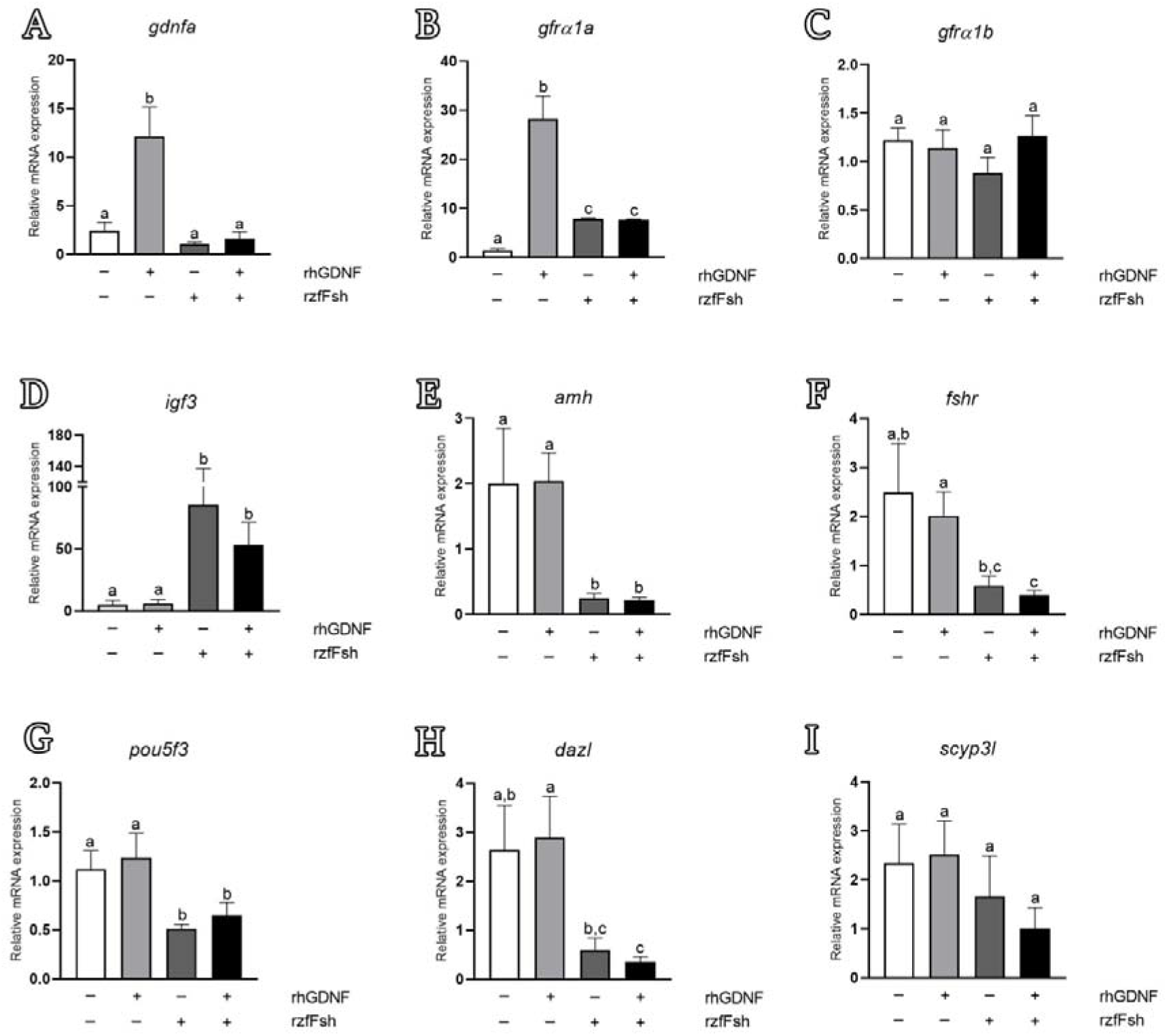
Relative mRNA levels of genes related to Gdnf signaling (*gdnfa*, *gfrα1a* and *gfrα1b*), Sertoli cell growth factors (*igf3* and *amh*), Fsh signaling (*fshr*) and germ cell markers (undifferentiated and differentiated spermatogonia - *pou5f3*; differentiated spermatogonia and preleptotene spermatocytes - *dazl*; and primary spermatocytes - *scyp3l*). Testicular explants were cultivated for 7 days with rhGDNF, rzfFsh or both (rhGDNF + rzfFsh). The relative mRNA levels were normalized with the *β-actin* levels. Bars represent the mean ± SEM (n = 20). Paired/unpaired t-test in which different letters denote significant differences (p < 0.05) among treatment conditions.

RT-qPCR analysis of zebrafish testis tissue *ex vivo* revealed that rhGDNF increased the transcript levels of *gdnfa* and *gfrα1a*, whereas *gfrα1b* mRNA levels remained unaltered when compared with basal condition levels (**Figure 8A-C**). The transcript abundance for the other genes (Sertoli cell growth factors, Fsh signaling and germ cell markers) did not change following rhGDNF treatment (**Figure 8D-I**). We further investigated whether rhGDNF could affect the Fsh-induced changes in testicular gene expression, since Fsh is considered the major endocrine player regulating zebrafish spermatogonial phase [56-57,66]. We first showed that Fsh did not modulate the transcript levels of *gdnfa*, *gfrα1a* or *gfrα1b* in the zebrafish testes (**Figure 8A-C**). However, Fsh was able to nullify the rhGDNF-increased *gdnfa* and *gfrα1a* mRNA levels following co-treatment (**Figure 8A, B**). With respect to Sertoli cell growth factors, we demonstrated that rhGDNF did not change the Fsh-mediated expression on *igf3* (**Figure 8D**) or *amh* mRNA levels (**Figure 8E**). As expected and in agreement with previous studies [67, 56], Fsh increased *igf3* mRNA levels (**Figure 8D**), and down-regulated *amh* transcripts (**Figure 8E**). The other evaluated genes were not responsive to either Fsh or to its co-treatment with rhGDNF (**Figure 8F-I**). Nevertheless, it is worth mentioning that transcript levels of *fshr*, *pou5f3* and *dazl* were significantly higher in the explants cultivated with rhGDNF than in those co-treated with rhGDNF + Fsh (**Figure 8F-H**).

### 3.7 *In silico* analysis of putative regulatory sequences upstream human *GDNF*, mouse *Gdnf* and zebrafish *gdnfa*

To support our expression analysis, we investigated the putative regulatory sequences upstream the transcriptional start site (TSS) of human *GDNF* (NM_000514.4), mouse *Gdnf* (NM_010275.3), and zebrafish *gdnfa* (NM_131732.2) (**Figure 9**). The *in silico* analysis showed three different types of cAMP response elements (CRE), several N-box and E-box motifs, one NF-kB binding site and a TATA-Box within the 2000bp upstream of human GDNF (**Figure 9, Table S2**). The upstream sequence of the *Gdnf* mouse gene showed similar regulatory binding sites as the human *GDNF* (**Figure 9, Table S2**). For zebrafish, we predicted a non-canonical TATA-Box, one CRE close to a GC-Box, one N-Box, four E-Box and two androgen receptor (AR) half binding site within 2000bp upstream of *gdnfa* (**Figure 9, Table S2**).

**Figure 9.**
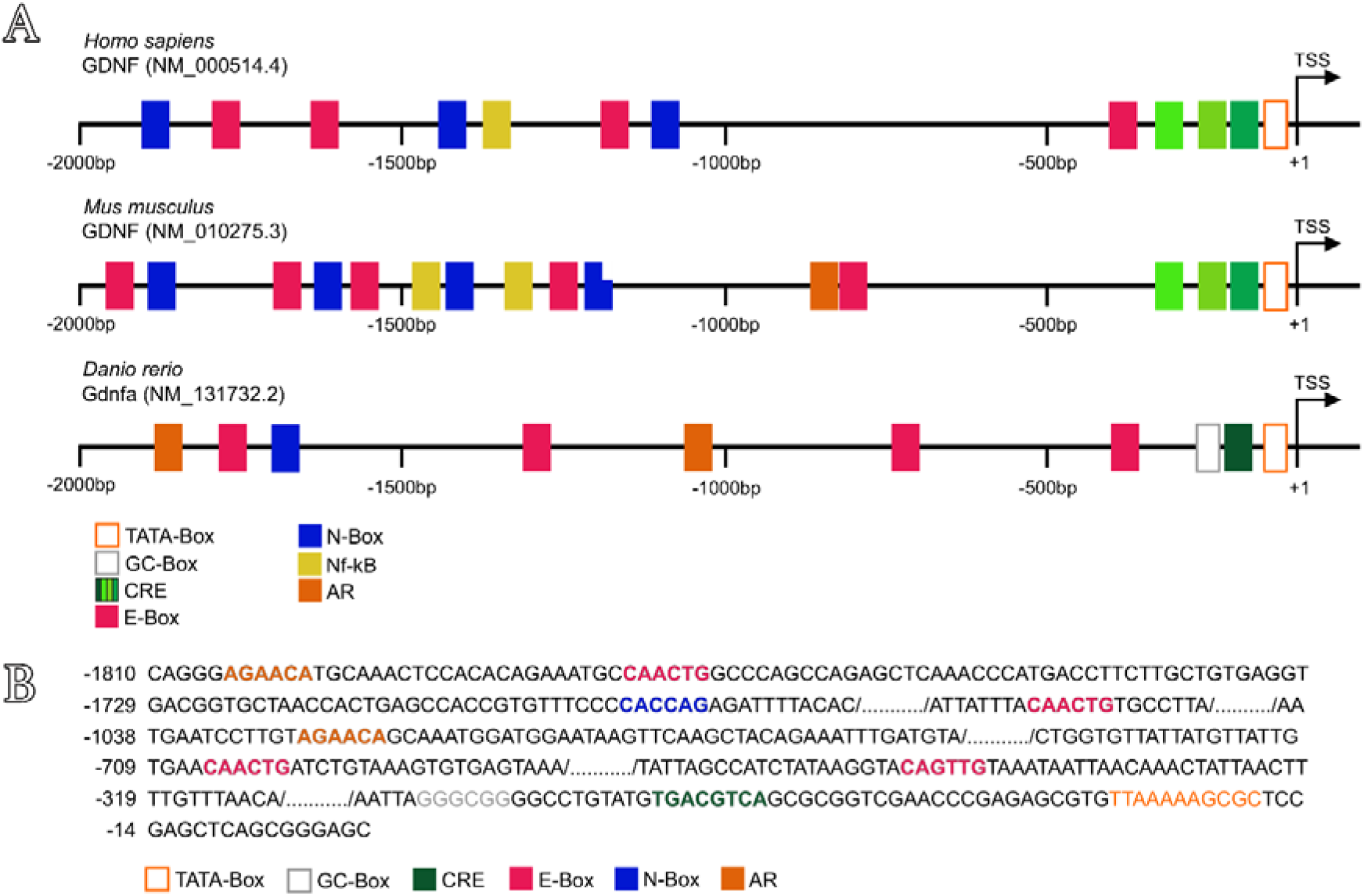
Predicted regulatory sequences upstream human *GDNF*, mouse *Gdnf* and zebrafish *gdnfa*. (A) The upstream region (2000bp) of human *GDNF* contains three different sequences of cAMP response elements (CRE), four E-box sequences, three N-box sequences and one Nf-kB binding site. The upstream region (2000bp) of mouse *Gdnf* contains a N-box/E-box-rich region at bp -1,300 to - 1,900 and additional E-boxes downstream, one androgen receptor binding site (AR), two Nf-kB binding sites and three different sequences of CRE close to the TSS (transcriptional start site). The upstream region (2000bp) of zebrafish *gdnfa* contains four E-box sequences and one N-box sequence, two AR half sequences and only one CRE close to a GC-box and the TSS. TSS is the transcription start site (position +1). (B) Sequence of putative binding sites upstream of zebrafish *gdnfa*. In the opened orange box is shown the TATA-box sequence, in gray the GC-box, in dark green the CRE, in pink the E-box, in blue the N-box, and in the orange closed box the AR half binding site.

## 4. Discussion

This study demonstrated the involvement of Gdnf/Gfrα1 signaling pathway in the regulation of zebrafish spermatogonial phase. Our first analysis identified two zebrafish paralogs for the Gfrα1 encoding gene, named as zebrafish *gfrα1a* and *gfrα1b*. The predicted amino acid sequence of zebrafish Gfrα1a and Gfrα1b revealed high identity to GFRα1 from other mammalian species investigated in this study (>60% and >57% sequence identity for Gfrα1a and Gfrα1b, respectively). Moreover, both paralogs have conserved domains and residues which are typical of the GFRα1 family members, such as three cysteine-rich domains (D1, D2, D3), 28 cysteine residues (plus 2 in the terminal region) and two triplets (MLF and RRR) [68–70]. Studies in mice using site-directed mutagenesis have shown that some of these conserved regions (e.g. two triplets - MLF and RRR - in the D2) are critical for Gfrα1 binding to Gdnf, activating the receptor complex and eliciting downstream signal transduction [68, 70]. This evidence suggested that theoretically both zebrafish Gfrα1a and Gfrα1b could bind and elicit a response to Gdnf/GDNF (e.g. rhGDNF). Moreover, in agreement with previous studies [71–72], phylogenetic analysis demonstrated that zebrafish Gfrα1a and 1b are clustered to other fish Gfrα1a and 1b, and the paralogs diverged and form two distinct sub-clades within the fish clade. Additional analysis of chromosome neighboring genes revealed that both zebrafish *gfrα1a*- and *gfrα1b*- containing regions are syntenic to human *GFR*A*1*- and rodents *Gfr*α*1*-containing regions. Altogether, this evidence confirmed that zebrafish *gfrα1a* and *gfrα1b* are duplicated genes that diverged from each other after the teleost-specific whole genome duplication. Around 320 million years ago, it is well established that the common ancestor of the teleosts experienced a third round of whole genome duplication [73–74]. This event was responsible to generate a large number of duplicated genes that could follow different evolutionary paths such as co-expression (both copies retain the ancestral function), non-functionalization (function loss or complete deletion of one copy), sub-functionalization (specialization of each copy, sub function partition), or neo-functionalization (acquisition of a novel function) [73–74]. In this study, we could not conclude the specific roles of Gfrα1a and Gfrα1b, although data on protein localization and expression analysis suggested that zebrafish Gfrα1a is the mammalian GFRα1 homologous copy (see discussion below). However, additional studies (e.g. specific knockout for each copy) are required to confirm this hypothesis, and to unravel the specific role of each Gfra1 paralog in the zebrafish spermatogenesis.

When evaluating the expression profiling of Gfrα1a and Gfrα1b, we found that both paralogs are expressed in the zebrafish testes, although *gfrα1a* transcripts were significantly more abundant than *gfrα1b*. Focusing our analysis in the zebrafish Gfrα1a, we developed a specific antibody to identify the cell types expressing Gfrα1a in the adult zebrafish testis. Our data revealed that Gfrα1a was detected in all types of zebrafish spermatogonia, although the staining pattern varied among the different generations of spermatogonia. Gfrα1a was mainly expressed in early types of spermatogonia (A_und_ and A_diff_), and its expression gradually decreased as spermatogonial clones became larger and more differentiated. Likewise, accumulative evidences have demonstrated that GFRA1 is a conserved marker for all types of undifferentiated spermatogonia in different mammalian species [22-29,75-76] and the frequency of GFRA1+ spermatogonia decreases as spermatogonia progress from A_s_ to Aal [76]. Similarly, in other fish species, mRNA or protein levels of Gfrα1a were found mainly in type A_und_ spermatogonia of dogfish (*Scyliorhinus canicula*) [30, 31], rainbow trout (*Oncorhynchus mykiss*) [32–34], medaka (*Oryzias latipes*) [35] and tilapia (*Oreochromis niloticus*) [78]. In rainbow trout, Nakajima et al [32] reported that *gfrα1* transcripts decreased throughout the spermatogonial development, and became undetectable in spermatids and spermatozoa. In medaka, Zhao et al [71] showed a moderate signal for *gfrα1a* and *gfrα1b* mRNA in spermatocytes, but no expression was found in spermatids and spermatozoa. Altogether these evidence are in agreement with our results, and support our hypothesis that Gndf-Gfrα1a signaling pathway is important for the regulation of zebrafish spermatogonial phase, but is not required for meiotic and post-meiotic phases. Strikingly, our study also detected the Gfrα1a protein among Sertoli cells associated with different types of germ cells. In rainbow trout, Maouche et al [72] demonstrated that *gfrα1a1* transcripts were mainly expressed in somatic testicular cells, while *gfrα1a2* was restricted to type A_und_ spermatogonia. To our knowledge, our study and the one in rainbow trout [72] were the first evidences to suggest that Gndf-Gfrα1a is not only involved in the control of spermatogonial phase, but also can modulate the functions of the cyst-forming Sertoli cells.

Investigation of the expression profiling of Gdnf ligands (Gdnfa and Gdnfb) revealed that *gdnfa* transcripts are predominantly more abundant than *gdnfb* (Ct = 28,54 for *gdnfa vs.* Ct = 31,04 for *gdnfb*), indicating that Gdnfa might be the main ligand in the zebrafish testis. Further *in situ* hybridization and RT-qPCR analysis demonstrated that *gdnfa* was mainly expressed in the zebrafish germ cells. No expression was observed in somatic testicular cells for *gdnfa*. In both analyses, we were not able to identify the specific germ cell types that express *gdnfa* in the zebrafish testis. Nakajima and collaborators [32], on the other hand, demonstrated in rainbow trout immature testes that *gdnf* mRNA and protein were expressed in type A_und_ spermatogonia. Moreover, the same authors showed that *gdnf* and *gfrα1* were co-expressed in germ cells, and their expression changed synchronously during the germ cell development [32]. Altogether, this evidence supports our findings that zebrafish Gdnfa is a germ cell derived factor that exerts both autocrine and paracrine functions on spermatogonia and Sertoli cells, respectively. Moreover, this data provides new insights on Gndf-Gfrα1a signaling pathway in fish when compared to mammals. In mammals, GDNF is secreted by testicular somatic cells (Sertoli cells [2,10,11], peritubular myoid cells [12, 13] and testicular endothelial cells [14]), acting only as a paracrine factor on GFRA1-expressing undifferentiated spermatogonia [22-29, 75-76]. This difference in the Gdnf-producing sites is likely related to events that took place after the teleost-specific whole genome duplication, such as non- and neo-functionalization of the Gdnf paralogs. In addition, these findings indicate that the common ancestor of Gdnf was expressed in the testicular somatic cells of fish and mammals, and the expression of Gdnf in the germ cells is an evolutionary novelty in the fish group. To support this hypothesis, identification of Gdnf paralogs and their cellular site expression in other vertebrate species, including fish, are necessary.

The biological roles of Gdnf on zebrafish spermatogenesis were assessed through employing a recombinant human GDNF (rhGDNF) in an *ex vivo* testis culture system previously established for zebrafish [53a]. There is strong evidence that human recombinant hormone (rhGDNF) can bind to zebrafish Gfrα1a and elicit a downstream signal transduction in the zebrafish testes. The first evidence is the predictive 3D model which examines the interaction sites between human GDNF and zebrafish Gfrα1a based on the binding interaction with human GFRA1. This analysis revealed structural similarities between zebrafish Gfrα1a and human GFRA1 (**Figure 6A, box2**), and higher identity of the structure formed at the binding sites between human GDNF and both receptors, human and zebrafish GFRA1/Gfrα1a, respectively (**Figure 6A, box2**). Moreover, this analysis also showed that most of the amino acid residues identified as crucial for ligand-receptor interaction are conserved in the zebrafish Gfrα1a, with exceptions for the residues Gly155 and Ile175, which were replaced by Glu and Thr, respectively. The predictive 3D model was also supported by Ramachandran plots which showed that 89.8% of the amino acid residues were in the most favorable regions, 7% of residues situated in allowed regions (∼2% expected), and 3.1% in the outlier regions. The second evidence is the sequence alignment demonstrating conserved regions between zebrafish Gdnfa (most abundant ligand) and rhGDNF, in particular, in the binding sites to human GFRA1 and zebrafish Gfrα1a. Lastly, the biological effects *per se* (e.g. proliferation and gene expression - see below) are the third evidence that rhGDNF not only can bind to zebrafish Gfrα1a, but also trans-activates the receptor complex to trigger molecular and cellular responses in the zebrafish testis.

With regards to biological functions, our results demonstrated that rhGDNF (100 ng/mL) increased the mitotic index of types A_und_ and A_diff_ spermatogonia when compared to basal condition. Consistently, histomorphometric analysis revealed that both types A spermatogonia (A_und_ and A_diff_) became more abundant, while type B significantly decreased following the rhGDNF treatment. Altogether, these results indicated that Gdnf not only stimulates proliferation of the most undifferentiated spermatogonia (A_und_ and A_diff_), but is also involved with blocking late differentiation into type B spermatogonia. Similar functions have been described for GDNF/Gdnf in mammalian and non-mammalian species. In mammalian species, particularly rodents, Gdnf promotes self-renewing proliferation of SSC [2]; see review in Parekh et al. [8] and Mäkelä and Hobbs [9]), although a recent study in mice has shown that Gdnf could be more associated with blocking differentiation rather than actively stimulating SSC proliferation [7]. In dogfish, rhGDNF promoted *in vitro* proliferation and long-term maintenance of spermatogonia with stem characteristics [31]. In medaka, Wei et al. [36] demonstrated that recombinant medaka Gdnfa and Gdnfb were involved in the proliferation and survival of medaka SSCs. Furthermore, the knockdown of medaka *gfrα1a* and *gfrα1b* subsequently confirmed that both receptors mediated the proliferation and survival of medaka SSCs [71]. In this study, Zhao et al [71] also showed that genes related with differentiation (e.g. *c-kit*) were up-regulated when lowering the expression of both receptors. Altogether, these evidences in different species sustain our conclusion that Gndf-Gfrα1 signaling pathway is associated with increasing/maintaining the pool of early types of spermatogonia (A_und_ and A_diff_) in the zebrafish testes through actively promoting their proliferation and also by inhibiting their differentiation. Moreover, as zebrafish Gdnfa and its receptor (Gfrα1a) are co-expressed, it is important to highlight that the above-mentioned effects are related to an autocrine loop of Gdnf on type A spermatogonia of zebrafish.

In this study, we also quantified Sertoli cell proliferation, reasoning that change in the proliferation of Sertoli cells associated with types A_und_ or A_diff_ spermatogonia would indicate creation of new niche space or supporting development of differentiating spermatogonial cysts, respectively [65]. In fish, in contrast to mammals, Sertoli cells are not terminally differentiated and continue to proliferate during spermatogenesis of adult males of different species, which also includes zebrafish [54, 75, 79]. Strikingly, our results demonstrated that Gndf promotes proliferation of Sertoli cells that are particularly associated with types A_und_ and A_diff_ spermatogonia which are also undergoing mitosis (BrdU-positive cells). This data indicates for the first time that a germ cell derived factor is involved in the creation of new spermatogenic cysts, i.e. new available niches, as well as supporting the development of early differentiating spermatogonial cysts. In the first case, as Gdnf stimulates proliferation of type A_und_, the newly formed, single spermatogonium must recruit its own Sertoli cells to form a new spermatogenic cyst. Therefore, it is reasonable that new Sertoli cells would be produced to create a niche into which the newly formed, single type A_und_ can be recruited or attracted (germ cell homing). Consistently, in mice, Gdnf has been shown to be important for germ stem cell homing as it acts as SSC chemotactic factor [80]. In the second case (supporting development of differentiating spermatogonial cysts), Gdnf-induced Sertoli proliferation would provide structural and nutritional support for the development of early differentiating spermatogonia. In both cases, Gdnf effects on Sertoli cells might be mediated directly through Gfrα1a which is also expressed in Sertoli cells of zebrafish. In agreement with our observation, it was reported in rodents that Gdnf also promoted the proliferation of immature Sertoli cells through its interaction with Gfrα1 and neural cell adhesion molecule (NCAM) both co-expressed in Sertoli cells [81–82]

We further evaluated whether Gdnf could modulate testicular gene expression or affect Fsh-induced gene expression in zebrafish explants. Previous studies have shown that Fsh is the major endocrine player regulating zebrafish spermatogonial development through targeting Sertoli and Leydig cells functions, such as sex steroid and growth factor production [49, 56, 66, 83, 84]. Our results showed that Gdnf positively modulates its own regulatory pathway (Gdnfa-Gfrα1a), increasing the transcript levels of both *gdnfa* and *gfrα1a* in the zebrafish testicular explants. This would be the first demonstration that a germ cell factor can affect the spermatogonial niche through an autocrine and paracrine loop. It seems that Gdnf signaling would enhance its own production and sensitivity to favor the creation of new spermatogonial niches (type A_und_ spermatogonia and Sertoli cells). Noteworthy, *gfrα1b* was not modulated by any treatment, which supports that zebrafish Gfrα1a could be the mammalian GFRα1 homologous form. Moreover, we showed that Fsh did not modulate *gdnfa* expression in zebrafish testicular explants. Similarly, Bellaiche et al [33] demonstrated that Fsh also did not modulate the expression of *gdnfb* in immature and early maturing rainbow trout testicular explants. This regulation in fish is different from the one reported in mammals, where Fsh has shown to stimulate the expression of *Gdnf* in the testes [84]. One possible explanation for this different regulation would be the distinct cellular sites expressing Gdnf in the mammalian and fish testes. In zebrafish, Gdnf is mainly secreted by germ cells, which are not the direct targets of Fsh, while in mammals, Gdnf is secreted by somatic cells, including Sertoli cells, which are known to express Fsh receptor. Additionally, to support our data, we performed *in silico* analysis within −2000 to +1 bp upstream of the zebrafish *gdnfa* gene to search cAMP response elements (CREs). As it is well known, Fsh activates the cAMP-dependent protein kinase A signaling pathway, resulting in phosphorylation of the cAMP response element-binding protein (CREB), which is required to transactivate several genes containing CREs [85]. Moreover, Lamberti and Vicini [60] demonstrated that three CRE binding sites in the murine *Gdnf* promoter are directly involved in the basal and cAMP-induced expression of *Gdnf* in murine Sertoli cells. In our *in silico* analysis, we demonstrated that zebrafish *gdnfa* promoter (−2000 to +1 bp) has less conserved DNA binding sites as comparable with human and mouse *GDNF*/*Gdnf* promoter. Moreover, our analysis showed only one CRE site near the zebrafish *gdnfa* transcription start site, instead of three CREs as reported in human and mouse. This difference in promoter region and quantity of CRE binding sites could be the reason that Fsh could not stimulate *gdnfa* expression in zebrafish.

The *GDNF*/*Gdnf* promoter region also contains several E-boxes and N-boxes that allow the binding of basic helix-loop-helix proteins with possible repressor activity on *GDNF*/*Gdnf* expression through Notch signaling [62]. Activation of the Notch receptor cleaves and releases the Notch intracellular domain (NICD) in the cytoplasm which migrates to the nucleus where it forms a transcriptional complex with the DNA-binding protein RBPJ (recombining binding protein suppressor of hairless) [88]. The canonical targets of RBPJ include the HES and HEY families of transcriptional repressors, which are basic helix-loop-helix proteins (bHLH) [89–91]. Transcriptional repressors of the HES family (HES1–7) bind to N-box promoter regions of their target genes, while repressors belonging to the HEY family (HEY1, HEY2, HEYL) bind to E-box promoter regions [90]. In zebrafish, it is known that Fsh stimulates Notch signaling [66]. Therefore, we speculate that Fsh nullified the Gdnf-increased *gdnfa* expression through Notch pathway and transcription repressors HES and HEY which would bind to N/E-boxes within the zebrafish *gdnfa* promoter region. However, functional studies on the zebrafish *gdnfa* promoter region are required to elucidate how Fsh and Gdnf regulate the expression of *gdnfa*.

In this study, we demonstrated that *gfrα1a* transcripts were up-regulated by Fsh, but not at the same intensity as in Gdnf treatment (3-fold increase as compared to Fsh). In immature rainbow trout, Bellaiche et al [33] also reported that *gfra1a* mRNA levels were increased following *in vitro* treatment with Fsh (100 ng/mL - same concentration as used in our work). Moreover, the same authors reported that testicular *gfra1a* levels increased towards the end of the reproductive cycle which coincides with the natural elevation of plasma Fsh levels in rainbow trout [33]. Therefore, different from mammalian species where Fsh up-regulated *GDNF*, we have evidence from two teleost species that Fsh modulates the Gdnf-Gfrα1 pathway through stimulating not the ligand, but the receptor (*gfra1a*) mRNA levels. However, there are some questions that remain unclear. The first question concerns whether the Fsh-induced expression of *gfra1a* is mediated by Sertoli, germ cells or both. In this work, we have demonstrated that *gfra1a* is expressed by Sertoli and germ cells, while the Fsh receptor is exclusively expressed by somatic cells (Sertoli and Leydig cells) [54]. Therefore, if Fsh-induced *gfra1a* expression is mediated by germ cells, this indicates that the regulation occurs indirectly through growth factors or androgens released by somatic cells (Sertoli and Leydig cells). Moreover, we cannot exclude that the increase of *gfra1a* could also be a consequence of the proliferation of spermatogonia or/and Sertoli cells stimulated by Fsh. Altogether, more studies are necessary to address the nature of Fsh regulation on *gfra1a* expression levels in the zebrafish testis. Although Gdnf or Fsh, independently, stimulated *gfra1a* mRNA levels in the zebrafish testis, we observed that co-treatment affected negatively the Gdnf-induced expression of *gfra1a* in the zebrafish testicular explants. This is also noted for other genes such as *pou5f3* and *dazl*, whose expression were higher in the Gdnf treatment as compared to co-treatment with Fsh. For *pou5f3*, a stem cell marker, this observation suggested that Gdnf could be more involved in the maintenance of stemness than increasing the number of stem cells in the zebrafish testis. On the contrary, Fsh would be more associated with proliferation towards differentiation, as *pou5f3* was significantly decreased following Fsh co-treatment. Therefore, our data indicate that the pro-differentiating effects of Fsh seemed to be more potent over the stem cell maintenance properties of Gdnf. On the other hand, at the level of differentiation, Gdnf decreased the Fsh effects on spermatogonial differentiation as the expression of *dazl*, a marker of late spermatogonial differentiation, was significantly down-regulated. Altogether, this observation suggests that Gdnf could promote stem cell maintenance through blocking spermatogonial differentiation. This conclusion is also supported by histomorphometrical data showing that Gdnf decreased the frequency of type B spermatogonia, and, somehow in agreement with the higher expression of Gfrα1a in type A_diff_ spermatogonia, which might be the one of the principal targets of Gdnf in the zebrafish testes.

As Gdnf is a member of TGF-β superfamily, its role on inhibiting spermatogonial differentiation is likely consistent with other TGF-β superfamily member’s role, such as Amh. Amh is a Sertoli cell growth factor which has been characterized as an inhibitor of spermatogonial differentiation in zebrafish [57, 67 92], see review in Adolfi et al, [93]. In this regard, we also examined whether Gdnf role could be modulated through Amh or by inhibiting Igf3, a pro-differentiation Sertoli cell growth factor (56-57, 83-84). Our data showed that rhGDNF did not modulate either *amh* or *igf3* mRNA levels in the zebrafish testicular explants. Therefore, Gdnf role on inhibiting zebrafish spermatogonial differentiation is not mediated by Amh or Igf3, and it could be either mediated directly on germ cells (autocrine) or indirectly through a different unknown growth factor released by somatic cells (paracrine).

**Figure 10** depicts our main findings regarding Gdnf actions on zebrafish testis. Gdnf is a germ cell growth factor that acts on type A spermatogonia and Sertoli cells in an autocrine and paracrine manner, respectively. Gdnf receptor, named as Gfrα1a, is expressed in type A spermatogonia (highly expressed in types A_und_ and A_diff_) and Sertoli cells. The main actions of Gdnf are: 1) creation of new available niches by stimulating proliferation of both type A_und_ spermatogonia and their surrounding Sertoli cells. In this context, we highlight that Gdnf stimulates proliferation of Sertoli cells which are associated with type A_und_ undergoing mitosis. As consequence, Gdnf increases the number of available niches and maintains the stemness pool in the zebrafish testes. 2) Gdnf also supports the development of differentiating spermatogonial cysts through proliferation of type A_diff_ and their surrounding Sertoli cells; and finally, it also 3) inhibits late spermatogonial differentiation as shown by the decrease of type B spermatogonia and down-regulation of *dazl* in the co-treatment with Fsh. Altogether our data indicates that although the autocrine and paracrine roles of Gdnf are evolutionary novelties in fish, Gdnf still exhibits similar/conserved functions as regards the mammalian GDNF. Our data showed that Gdnf is not increasing the number of SSCs in the testis, but rather is responsible for maintaining the spermatogonial stemness in the zebrafish testes by tightly regulating the processes of creation of new available niches, supporting development of early differentiating spermatogonial cysts and inhibiting late spermatogonial differentiation.

**Figure 10.**
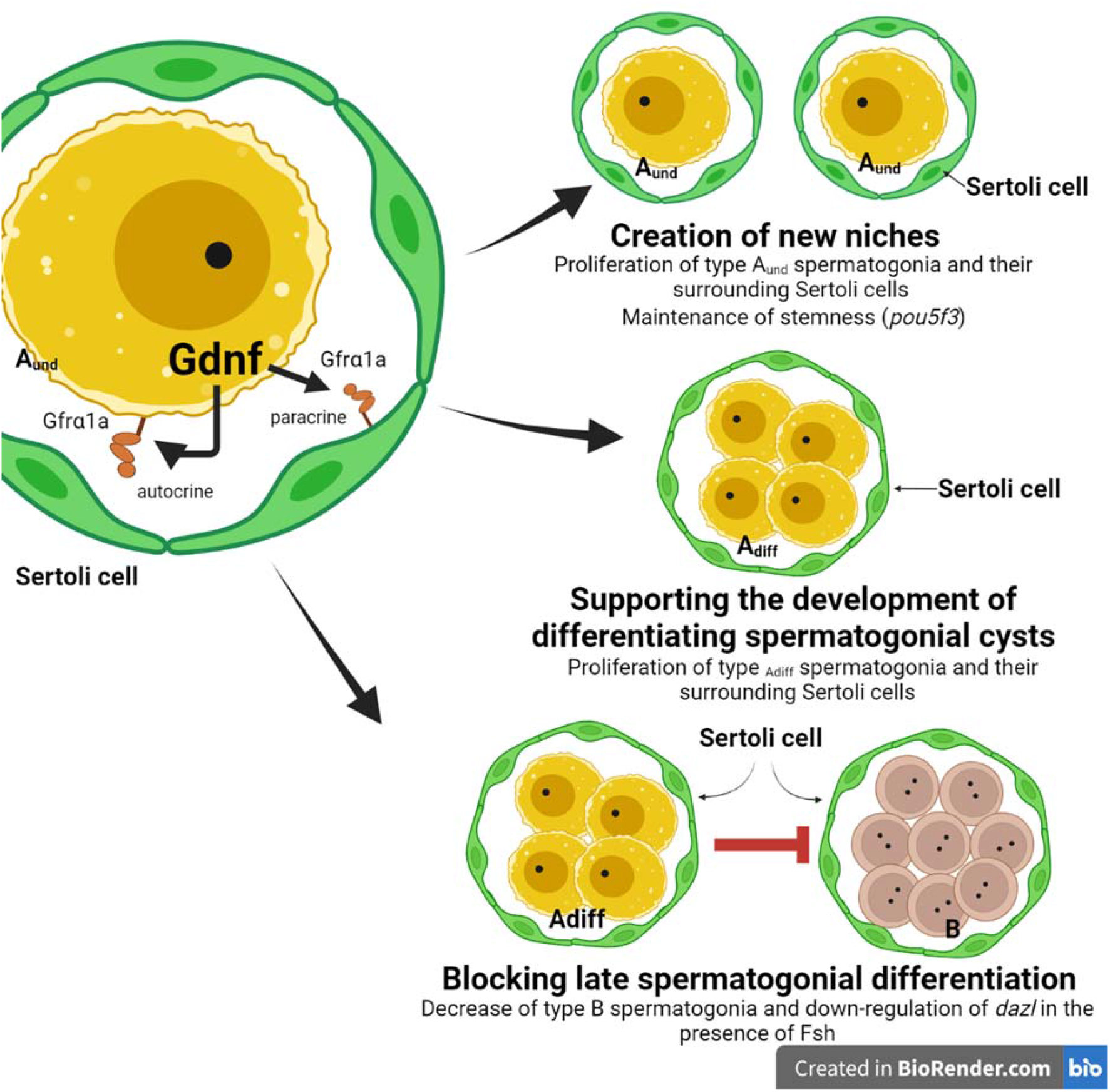
Summarizing effects of Gdnf in the zebrafish spermatogonial niche. Gdnf is a germ cell growth factor which acts on type A spermatogonia and their surrounding Sertoli cells in an autocrine and paracrine manner, respectively. Gdnf receptor, named as Gfrα1a, is expressed in type A spermatogonia (early spermatogonia, with higher expression in types Aund and Adiff) and Sertoli cells. The main actions of Gdnf are: 1) creation of new available niches; 2) supporting the development of early differentiating spermatogonial cysts; and 3) blocking late spermatogonial differentiation.

## Acknowledgments

This research was supported by São Paulo Research Foundation (FAPESP) (2016/12101-4/ 2017/08274-3 granted to L.B.D.; 2014/07620–7 and 2020/03569-8 - granted to R.H.N.) and financed in part by the Coordenação de Aperfeiçoamento de Pessoal de Nível Superior – Brasil (CAPES) – Finance Code 001” (granted to L.D.B.). R.H.N and G.M. are granted with productivity scholarships from Brazilian National Council for Scientific and Technological Development (CNPq) (proc. n. 305808/2020-6 and 307743/2018–7, respectively).

## Authors Contribution

Conceived and designed the experiments: RHN, LBD, AJB, ERMM. Performed the experiments: LBD, AJB, ERMM, RTN, BM, JMBR, IFR, MRS, ATN, DFC. Analyzed the data: LBD, AJB, BM, RTN. Contributed reagents/materials/analysis tools: RTN, RHN. Wrote the manuscript: LBD, GM, CS, RHN.

## Conflicts of Interest

The authors declare no conflict of interest.

## Supplemental material

### 1. Material and Methods

#### 1.1 gdnfa expression in zebrafish adult testis

For *in situ* hybridization, a zebrafish *gdnfa*-specific PCR product was generated with primers *gdnfa-ish*-Fw and *gdnfa-ish*-Rv (Table 1). The ∼160 bp PCR product was gel purified, and served as a template for digoxigenin (DIG)-labelled cRNA probe synthesis using the RNA labeling (Roche) kit. Gonads were fixed in 4% paraformaldehyde (PFA) in PBS at 4°C for 2 hours. The protocol used for whole mount (WISH) and in situ hybridization (paraffin embedded) were performed with adaptations, as described previously [1]. Detection of hybridization signal was done with HNPP Fluorescent Detection Set (Roche). Nuclei counter-staining was performed with DAPI (Sigma) (1:10000) diluted in PBS (Phosphate Buffered Saline pH 7.4, sterile-filtered).

### 2. Results

#### 2.1 Morphological characteristics and in situ hybridization for gdnfa in adult zebrafish testis

**Supplemental Figure 1.**
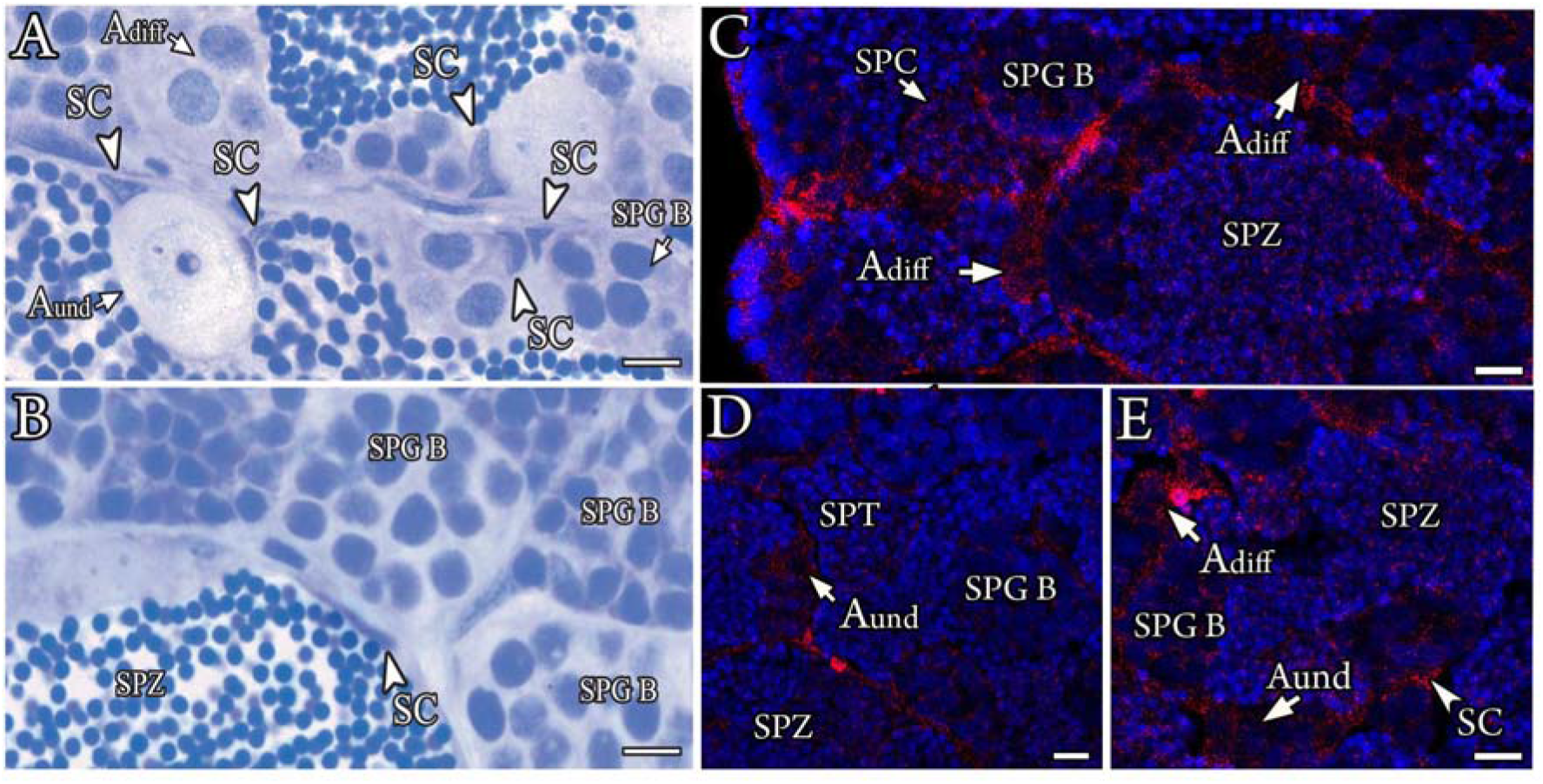
Morphological characteristics and *in situ* hybridization for *gdnfa* in adult zebrafish testis. A-B. Adult zebrafish testis section showing different germ cysts (arrow) and Sertoli cells (arrowhead). Bars-5µM C-E. Detection of *gdnfa* mRNA by *in situ* hybridization. Bars - 10 µM. Aund – type A undifferentiated spermatogonia, Adiff type A differentiated spermatogonia, SPG B – type B spermatogonia, SPC – spermatocytes, SPT – spermatids, SPZ – spermatozoa and SC - Sertoli cell

#### 2.2 Control sections using either preadsorbed antibody with the corresponding peptide or omitting the antibody

**Supplemental Figure 2.**
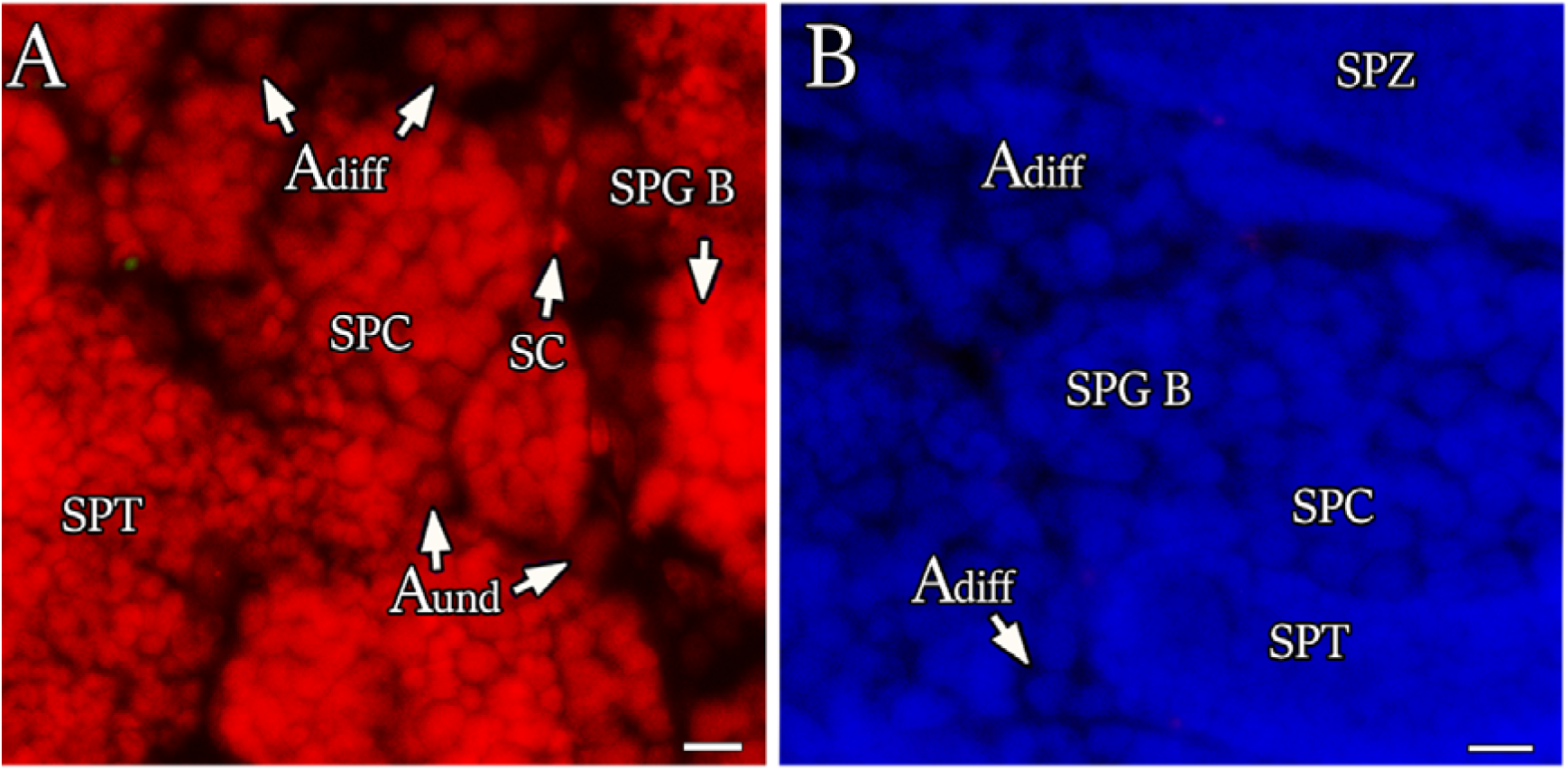
Control sections of immunofluorescence of cellular localization of Gfrα1a protein in zebrafish testis (green - A; red – B) using either preadsorbed antibody with the corresponding peptide or omitting the antibody confirming the antibody specificity. Bars - 10 µM. Aund – type A undifferentiated spermatogonia, Adiff type A differentiated spermatogonia, SPG B – type B spermatogonia, SPC – spermatocytes, SPT – spermatids, SPZ – spermatozoa and SC - Sertoli cell.

#### 2.3 Detailed interaction between rhGDNF and the zebrafish receptor Gfrα1a

**Supplemental Figure 3.**
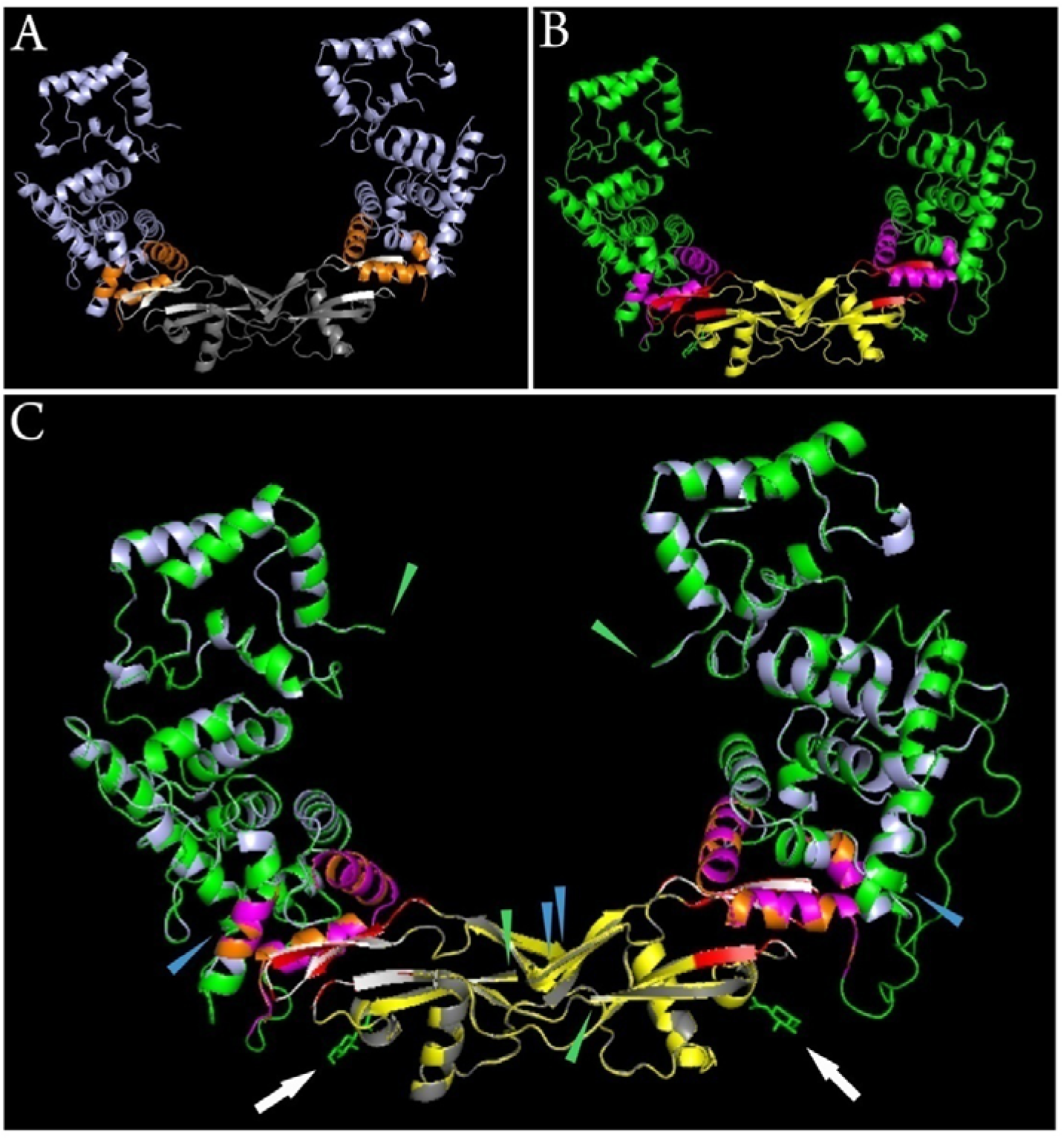
Predicted protein complex models of *Danio rerio* Gfrα1 and rhGNDF (hetero-2-2-mer). In A, the template is chosen according to results performed by SWISS-MODEL (swissmodel.expasy.org). Blue arrowheads show the Gfrα1 protein structure and in orange the ligand sites bind to rhGNDF protein. Dark grey represents the rhGNDF proteins and light grey the ligand sites bind to Gfrα1. In B, the 3D model of the Gfrα1-Gndf protein structure. In green, the Gfrα1 protein structure and in purple the ligand sites bind to Gndf protein. Yellow represents the rhGNDF proteins and in red the ligand sites that bind to Gfra1a. Arrows indicating two ligands of N-Acetyl-D-glucosamine. In C, we merged the template and the model of the hetero-2-2-mer showing the similarity of the structures and the identity of the structure formed at the binding sites between Gfrα1 and rhGNDF proteins (orange-purple and light grey-red). Arrows indicating two ligands of the ligand N-Acetyl-D-Glucosamine.

### Video 1 here

**Video 1.** Video of the detailed interaction between the rhGDNF and its possible ortholog receptor in zebrafish adult testis, Gfrα1a. Despite the evolutionary distance between both receptor and ligand, the ligand is still able to respond to the recombinante human GDNF.

**Table S1.**
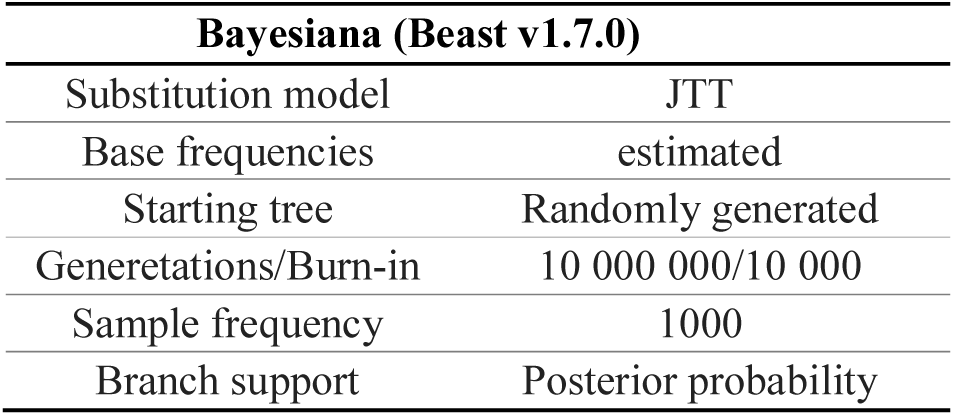
Parameters set to reconstruct the phylogeny.

**Table S2.**
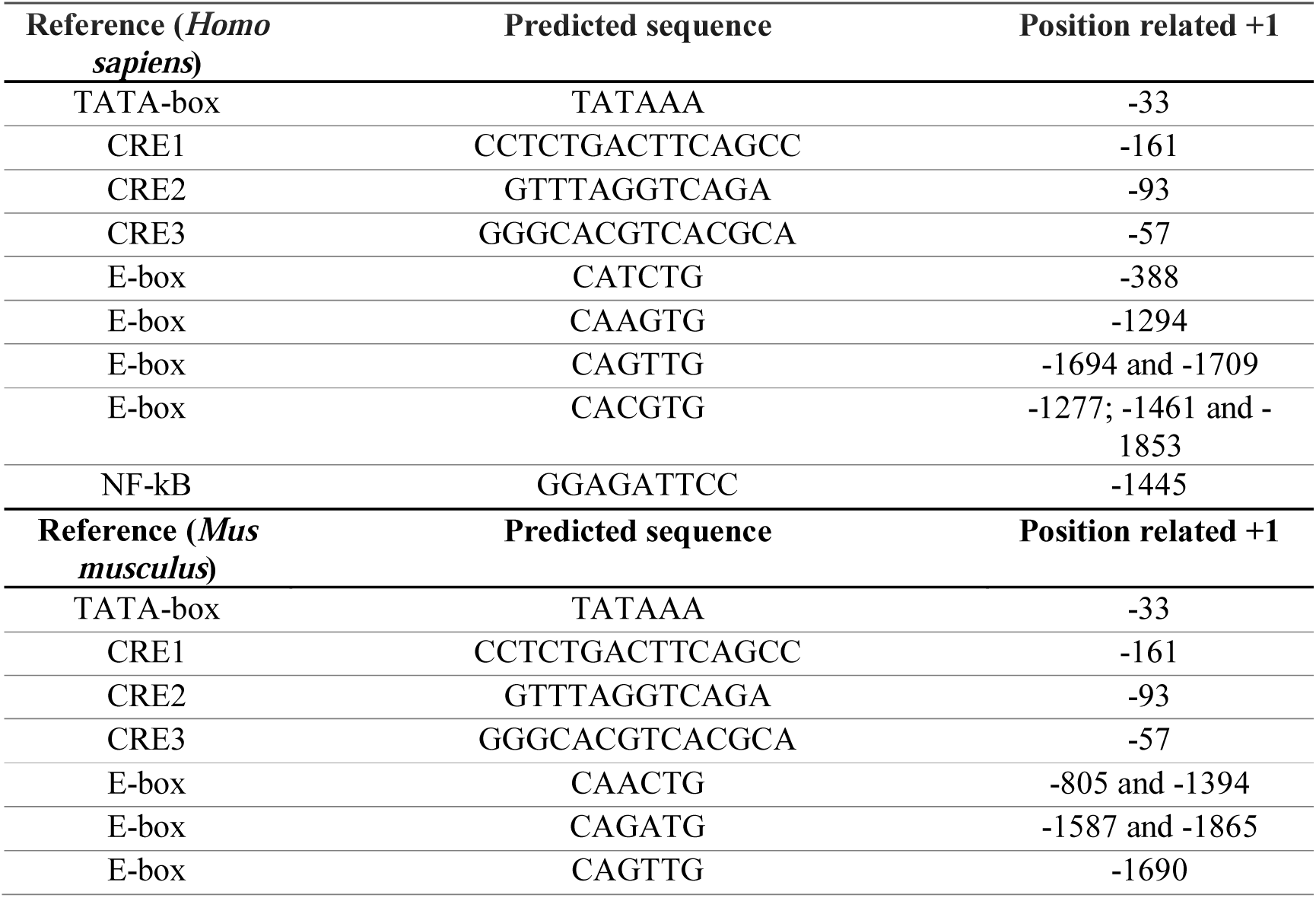
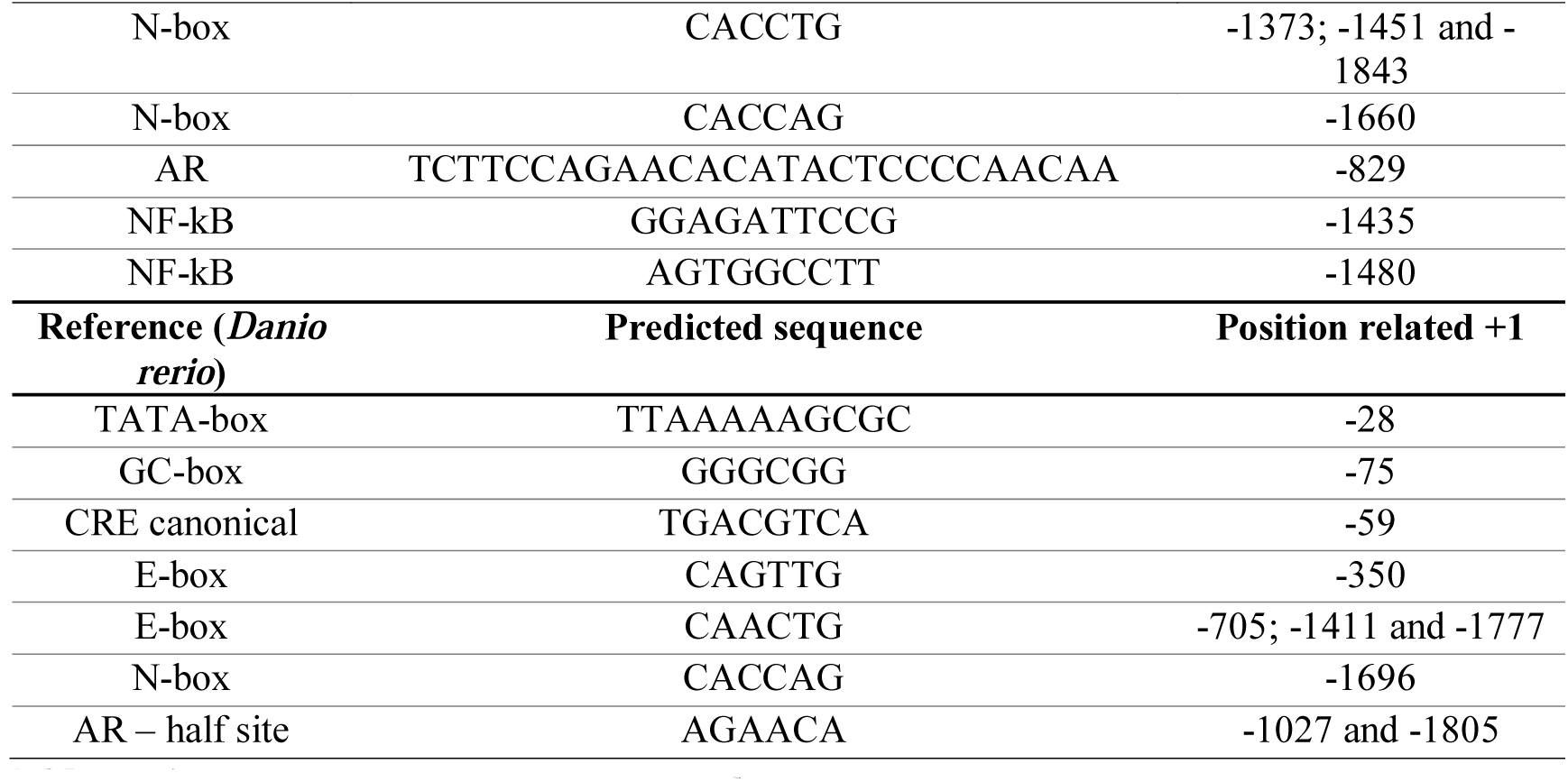
Predicted regulatory binding sites of the GDNF promoter in *Homo sapiens*, *Mus musculus* and *Danio rerio*.

## Notes

### Competing Interest Statement

The authors have declared no competing interest.

